# DNA metabarcoding diet analysis in ruminants is quantitative and integrates feeding over several weeks

**DOI:** 10.1101/2024.02.01.577814

**Authors:** Stefaniya Kamenova, Pernille Meyer, Anne Krag Brysting, Leo Rescia, Lars P. Folkow, Monica Alterskjær Sundset, Éric Coissac, Galina Gussarova

**Author notes:** Equal contribution (Éric Coissac, Galina Gussarova).

## Abstract

Dietary DNA metabarcoding is an established method, especially useful for resolving the diverse diets of large mammalian herbivores (LMH). However, despite longstanding research interest on the topic, we still lack unequivocal evidence on the potential of DNA metabarcoding to reflect proportions of ingested dietary plants in LMH. One major aspect to consider is the time window during which ingested diet remains detectable in faecal samples. This parameter is currently unknown for LMH, thus potentially hindering the scope of ecological conclusions. Another unknown factor is quantitative performance, i.e. the extent to which the amount of ingested biomass can be assessed based on sequence read abundances. We assessed DNA metabarcoding quantitative performance and DNA half-life detectability for plants with different digestibilities in a controlled feeding experiment with three female Eurasian tundra reindeer (*Rangifer tarandus tarandus*). Reindeer were fed birch twigs (*Betula pubescens*) and increasing biomass of lichen (mainly *Cladonia stellaris*). Relative read abundance positively correlated with ingested lichen biomass, suggesting potential for deriving dietary proportions in free-ranging reindeer on natural pastures. Dietary DNA was consistently detected within a few hours upon ingestion, with a mean half-life detectability of 30 and 14 hours for birch and lichen, respectively. However, dietary DNA remained detectable in faeces for at least 26 days post-feeding, indicating that a single faecal sample can provide an unsuspectedly integrative estimate of diet in ruminants. Together, our findings provide novel empirical validation of DNA metabarcoding as a tool for diet analysis in LMH.

## 1 Introduction

DNA metabarcoding diet analysis from faecal samples is a highly sensitive, cost- and time-efficient method for assessing animal diets (Pompanon et al. 2012; Ando et al. 2020). The method has encountered particular success with ecologists focusing on large mammalian herbivores (LMH) (e.g. Pansu et al. 2019; Jorns et al. 2020; Spitzer et al. 2021; Mitchell et al. 2022). LMH typically display highly diverse and dynamic diets, difficult to track with time-consuming approaches such as microhistology (e.g. Garnick et al. 2018), which is also subject to observer bias, while food items leaving no, or few visible residues can be entirely missed from the analysis (Soininen et al. 2009; Nichols et al. 2016). Although such challenges can be avoided by using DNA metabarcoding, this approach has other limitations, many of which remain largely unassessed in the case of LMH (but see Scasta et al. 2019). Overall, there is still a considerable knowledge gap regarding the process of plant DNA digestion in large herbivores, and, by consequence, our ability to robustly interpret DNA metabarcoding diet datasets is still limited.

One important knowledge gap is the temporal window of diets that may be captured by DNA metabarcoding of faecal samples. It is often assumed that DNA metabarcoding provides a rather short temporal snapshot of individual diets, due to the negative exponential decay of dietary DNA (Deagle et al. 2006; Weber & Lundgren 2009; Paula et al. 2014). This also coincides with estimates about food particles mean retention times, which in wild ungulates typically approximate 24 to 48 hours (Barboza et al. 2006; Steuer et al. 2011; Picard et al. 2015). LMH, especially ruminants, possess complex multi-chambered digestive systems, functioning in partnership with microbial symbionts for the breakup of recalcitrant carbohydrates from the plant diet (Hungate 1975; Kay et al. 1980; Dehority 2003). Ruminants also chew cud in order to optimise digestion. How this digestive complexity impacts the DNA digestibility and temporal decay of DNA from the food items is unclear. Based on the identification of a greater number of plant taxa with DNA metabarcoding in the rumen content from fallow deer (*Cervus dama*) and roe deer (*Capreolus capreolus*), Nichols et al. (2016) propose that DNA from the food material might be detectable for longer periods compared to the plant cuticles targeted with microhistological analysis. However, we still lack more precise temporal estimates of DNA half-life detectability (T_1/2_) - the post-feeding time required to reduce by half the amount of dietary DNA through digestion (Greenstone et al. 2014). This is important because food items varying in their digestibility could display differences in DNA decay dynamics, and thus detectability, which in turn could be misinterpreted as differences in dietary preferences. Earlier studies, based on diagnostic PCR in invertebrate predators, have already pointed out the risks of misinterpreting differences in T_1/2_ among different prey items as differences in consumer’s feeding rates or diet preferences (Chen et al. 2000; Greenstone et al. 2007; von Berg et al. 2008). LMH can typically feed upon a broad range of plant species, varying in their life-history traits and phenological stages, and thereby simultaneously ingest food items with very different digestibility characteristics. Thus, differences in digestibility among food items within the gut might impact their detectability, and by consequence, their relative proportions in the faeces. On the other hand, the occurrence of food items with differential DNA retention times might potentially result in the simultaneous detection of the partial remains of recent and older meals within individual faeces.

Quantitative performance, i.e. the extent to which sequence read abundances correlate with species numerical abundances or biomass, is another important aspect of dietary DNA metabarcoding for which we currently lack convincing data. Factors such as differences in PCR amplification efficiency, tissue cell density, interspecific variation in gene copy number, as well as differential digestibility among food items have all been shown to bias the proportion of recovered DNA sequences (Deagle & Tollit 2007; Deagle et al. 2010; Pompanon et al. 2012; Deagle et al. 2013; Neby et al. 2021; Stapleton et al. 2022). However, studies specifically focusing on LMH seem to suggest much higher congruence between DNA relative read abundance and ingested plant biomass (Willerslev et al. 2014), or between DNA relative read abundance and dietary proportions estimated with other methods - the weighed amount of plant fragments (Nichols et al. 2016) or the proportional consumption of C^4^ plants estimated from stable isotopes (Kartzinel et al. 2015). But the number of studies remain modest and experimental assessment of the relationship between DNA relative read abundance and food proportions in LMH is still missing.

The reindeer is a keystone species within the Arctic tundra ecosystem, sustaining local food webs as well as the livelihoods of numerous indigenous peoples (Turi 2002; Jernsletten & Klokov 2002; Müller-Wille et al. 2006). The complexity of the feeding behaviour displayed by reindeer makes them a highly relevant model in our effort to better understand the DNA digestive process in LMH. Reindeer are ruminant and intermediate mixed feeders highly adapted to the contrasted Arctic seasonality with changes in chemistry, digestibility, and availability of the pasture plants (Hofmann 1989; Hofmann, 2011; Mathiesen et al. 1999; Mathiesen et al. 2000a; Mathiesen et al. 2000b; Storeheier et al. 2003). Reindeer also rely on a unique and complex symbiotic microbiome of anaerobic bacteria, archaea, fungi and ciliates within their reticulo-rumen and distal fermentation chambers (caecum and proximal colon) for the digestion and utilisation of their heterogenous diet of arctic plants and lichen (Sundset et al. 2007; Sundset et al. 2009; Pope et al. 2012; Sundset et al. 2013; Salgado-Flores et al. 2016).

Here, we assess DNA metabarcoding quantitative performance and DNA half-life detectability using a controlled feeding experiment with three female reindeer (*Rangifer tarandus tarandus*) at the Arctic University of Norway in Tromsø, Norway (Meyer 2019). The three animals were fed birch twigs (*Betula pubescens*) and increasing amounts of lichen (mainly *Cladonia stellaris*). Both food items are palatable to the reindeer but have contrasting digestibility. Typically, winter birch twigs can be high in phenolic compounds (6% of dry matter - DM), which have been shown to inhibit digestibility (Palo 1985; Sunnerheim et al. 1988). Birch twigs also contain large amounts of lignin (24.7 % of DM) compared to other plants eaten by the reindeer. Lignin has an average *in vitro* digestibility of only 20.8% when using a ruminal inoculum from free-ranging reindeer (Storeheier et al. 2002a). For comparison, *in vitro* digestibility of whole thalli of the lichen species *Cladonia stellaris* was 55.7% DM, while as high as 77.2% DM for whole thalli from the lichen *Cetraria islandica* (Storeheier et al. 2002b). *Cladonia* lichens synthesise and accumulate phenolic compounds such as usnic acid, presumably representing a protective measure against grazing (Ingólfsdóttir 2002). However, the rumen microbiome of reindeer has evolved mechanisms to tolerate and detoxify these secondary compounds, rendering lichens a highly digestible food resource to reindeer in winter (Palo 1993; Storeheier et al. 2002b; Sundset et al. 2008; Sundset et al. 2010).

We hypothesize that (i) DNA relative read abundance is positively correlated with the ingested biomass of lichen, (ii) DNA half-life detectability of both birch and lichens would fall within the time window of 24-48 hours, but that the detectability of the more digestible lichen would decay significantly faster compared to birch.

## 2 Materials and methods

### 2.1 Experimental setup

The feeding experiment was carried out in the period January - February 2018 (Figure 1) at the Department of Arctic and Marine Biology, UiT - The Arctic University of Norway in Tromsø. Three 7-year-old pregnant female reindeer were selected for the experiment (animals 9/10, 10/10 and 12/10). At this period of year, the animals were kept in large outdoor enclosures where they were fed *ad libitum* pelleted feed (FK Reinfôr, Felleskjøpet, Norway). The composition of the pelleted feed was obtained in advance from the supplier (Table S1). The reindeer could also freely feed on the range of plants, mosses and lichens present within their outdoor enclosure. During the experiment, the reindeer were placed at a natural daylight cycle in smaller outdoor fan-shaped corridor enclosures, which were covered with heated tarmac to prevent snow and ice accumulation. They were accustomed to the experimental setting for a month prior to the experiment. Just before the start of the experiment, enclosures were thoroughly cleaned with a high-pressure water washer, and then sterilised using a propane burner aiming at removing all DNA traces from urine, faeces, food, and vegetation that might have been present. Animals were re-introduced into the corridor enclosures on January 19 and did not had access to outdoor pastures until February 18. During the course of the experiment, all faecal matter was removed after each sampling visit, while enclosures were cleaned once a day with a high-pressure water washer in order to eliminate faecal and urine residues. Water and feed were provided *ad libitum* in separate plastic containers, which were all thoroughly cleaned using 4% bleach solution at the start of the experiment. The quantity of feed provided, and all uneaten leftovers were recorded every day (Table S2). Since the experiment was completely non-invasive and had no conceivable consequences to either the health or physiology of the research animals, no specific authorised permit was required. The experiment was approved locally (permit # AAB-09).

**Figure 1.**
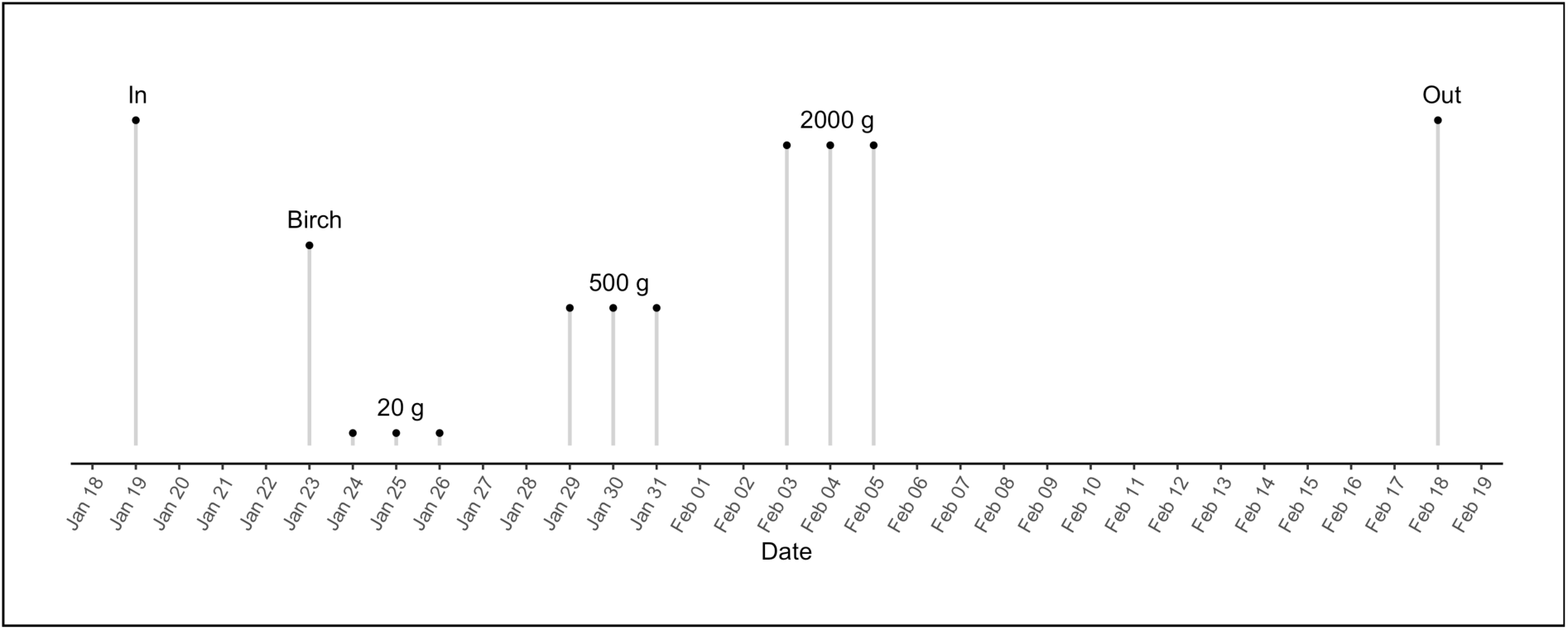
The timeline of the feeding experiment including the dates for moving the reindeer in and out of the experimental enclosures, and the dates when they were fed birch (*Betula pubescens*) and increasing biomass (20, 500 and 2,000 grams) of lichen (mostly *Cladonia stellaris*).

### 2.2 Feeding

Birch twigs (*Betula pubescens*) were collected in proximity to the reindeer large outdoor enclosures at UiT, while lichen (mainly *Cladonia stellaris*) was collected from different mountain regions in southern Norway. On January 23 in the evening, in addition to their daily ration of 2,000 grams of pelleted feed, reindeer were offered 50 g of birch twigs for two hours (Figure 1). After feeding, food containers were removed for cleaning and any birch leftovers recorded and removed from the corridor enclosures. On January 24, 20 g of wet lichen biomass was offered to each reindeer (Figure 1). Reindeer were let feeding for two hours before removing food containers for cleaning and lichen leftovers recorded and removed. The same procedure of lichen feeding was repeated for two more days. Following the same protocol, and within a gap interval of 48 hours every time, we kept introducing increasing amounts of lichen into the reindeer diet, respectively 500 and 2,000 g of wet lichen biomass offered to each individual (Figure 1). After introducing the last batch of lichen (February 5) and until the end of the experiment (February 18 – i.e. lasting ∼4 weeks), reindeer were exclusively fed pelleted food in the corridor enclosures.

### 2.3 Sample collection

Reference faecal samples were collected from reindeer (available only for 9/10 and 12/10) before the start of the experiment, on January 22, which was considered as Day 0. During the first 10 days after feeding birch, all faeces were systematically collected by monitoring each reindeer every 2 hours during daytime and every 3 hours after sunset. Night checks were carried out using infrared light to avoid disrupting the natural circadian rhythm of the animals during the polar night. After Day 11, faeces sampling was carried out every 3 hours, and during daytime only. The sampling effort was further reduced to one collection per day after Day 23. Faecal samples were also collected once or twice a week after the end of the experiment, after the reindeer had been returned to their outdoor enclosures with natural pastures (cf Table S3). All faecal pellets were collected directly from the ground in the corridor enclosures using disposable gloves, and immediately placed at -20°C into clean, pre-labelled zip-lock bags. Exact time of collection was systematically recorded. All excess pellets were systematically removed to minimise possible cross-contaminations among samples.

### 2.4 DNA metabarcoding diet analysis

We selected a total of 234 faecal time-points (representing 47% of all the collected faecal samples) for further analysis (Table S4). Samples were selected by dividing each sampling day into four time-intervals (00:00-06:00, 06:00-12:00, 12:00-18:00 and 18:00-24:00), and only selecting the last faecal sample of each time interval. Defrosted faecal pellets were thoroughly homogenised within the zip-lock bags and subsampled for DNA extraction. DNA extractions were carried out with the DNeasy PowerSoil kit (Qiagen, Germany) according to the manufacturer’s instructions. For covering the full range of reindeer diet, we used a combination of the *trn*L *Sper01* primers targeting the P6 loop in seed plants (Spermatophyta) (Taberlet et al. 2007), and the 18S *Euka02* primers targeting eukaryotes (Guardiola et al. 2015), see full details in Table S5. Further details about the PCR procedures can be found in Appendix 1. Libraries were prepared using the KAPA HyperPlus kit (Kapa Biosystems, USA), and sequenced on a HiSeq 4000 machine (Illumina, USA) following manufacturer’s instructions at the Norwegian Sequencing Centre (https://www.sequencing.uio.no). Bioinformatic analyses were carried out using the Norwegian high performance computing cluster Saga (https://www.sigma2.no) and the OBITools program (Boyer et al. 2016). Bioinformatic steps are described in Appendix 2.

### 2.7 Data curation

Datasets were imported in R v. 4.2.3 (R Core Team, 2016). Data curation was carried out according to Zinger et al. (2019) and filtering thresholds were based on the *ad hoc* examination of the sequence data. Full details are reported in Appendix 3 as well as in the archived analysis scripts files (https://renin-project.github.io/Feeding-Experiment/). After further inspection of the curated datasets, we retained only (i) MOTUs with 100% best-identity match to the reference database for *Sper01*, and 95% for *Euka02*, (ii) only MOTUs within the taxonomic range covered with each primer set (i.e. Spermatophyta for *Sper01*; Spermatophyta and Lecanoromycetidae, the subclass of lichen-forming fungi, for *Euka02*). Sequence read abundance was normalised by computing RRA using the *decostand* function from the R package “vegan”. The final *Sper01* dataset resulted in 45 plant MOTUs distributed among 217 reindeer faecal samples passing all quality criteria, while the final *Euka02* dataset resulted in 44 MOTUs distributed across 217 samples.

### 2.8 Statistical analyses

#### From RRA to DNA amount

DNA metabarcoding can provide quantitative information about dietary proportions based on RRAs of each MOTU. By definition, the sum of RRAs of all MOTUs in a given sample equals one, meaning that one degree of freedom is lost. Consequently, if the proportion of a dietary item increases (e.g. due to increased intake), RRAs of all other dietary items will automatically decrease. For the same reason, if the intake of a dietary item doubles, the RRA of that item will increase, but by a factor less than two. This effect is particularly marked when the variation in the quantity of an item is large, and the diversity of the diet is low. In our experiment, reindeer diet consisted of two parts: (i) a constant input of pelleted feed, supplied in the same quantity throughout the experiment, (ii) a variable part made up of birch or lichen, added punctually for the need of the experiment. According to the principle explained above, the addition of birch or lichen will reduce RRAs of plants composing the pelleted feed. To account for this effect and to get a better insight into the kinetics of birch and lichen DNA half-life detectability, RRAs of these two items, named respectively *RRA_birch_* and *RRA_lichen_,* were divided by the sum of the RRA of two selected plant MOTUs (Polygonaceae with *Sper01*, Brassicaceae with *Euka02*, cf Figures S1, S2), corresponding to the pelleted feed (*RRA_pellets_*) and multiplied by the mass of pelleted feed ingested the day before (*F_pellets_*), to account for the delay between the ingestion and the arrival of DNA in the faeces. The corrected value for the amounts of birch and lichen thus corresponds to *Q_birch_ = RRA_birch_ / RRA_pellets_ x F_pellets_* and *Q_lichen_ = RRA_lichen_ / RRA_pellets_ x F_pellets_*, respectively, and is expressed in an arbitrary unit of DNA amount, which was not standardized across animals.

#### Relationship between DNA amount (Q_item_) and biomass of ingested food

We expected that Q_item_ correlates with the biomass of birch or lichen ingested during the experiment. To test this assumption, lichen biomass (*F_lichen_*) fed to the reindeer was compared to the maximum amount of lichen DNA - *max*(*Q_lichen_*) - detected in reindeer faeces after each of the lichen meals using a linear model. *max*(*Q_lichen_*) is measured as the median of *Q_lichen_* observed between 12 and 60 hours after lichen feeding. This corresponds to the time intervals 36–84, 156–204, and 276–324 hours after the birch feeding for each of the three lichen biomasses fed to the reindeer (i.e. 20, 500, and 2,000 grams, respectively). To standardize *max*(*Q_lichen_*), expressed in arbitrary units depending on the animal, values for each animal were divided by *summax*(*Q_lichen_*) - the sum of the *max*(*Q_lichen_*) for the three lichen biomasses for each animal. The *lm* function in R was used to estimate the parameters of the model:

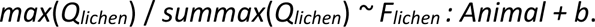

Significance for the three slopes (one for each animal) was assessed using a Student’s t-test and the global adjusted coefficient of variation R^2^_adj_ of the model was obtained from the model summary. To check the validity of the tests, the normality of the residuals of the model was tested using a Shapiro-Wilk test (Royston 1982).

#### Estimation of DNA half-life detectability

Typically, dietary DNA decay follows a negative exponential function (Deagle *et al*. 2006; Weber & Lundgren 2009; Paula *et al*. 2014). Therefore, DNA half-life detectability – *T_1/2_*_item_ (*T*_1/2birch_ or *T*_1/2lichen_) – can be calculated from the slope *a_item_* of a linear model:

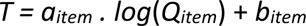

where *T* is the sampling time and *Q_item_* is the amount of dietary DNA (i.e. *Q_birch_* or *Q_lichen_*), and therefore expressed as:

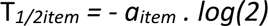

The overall time frame for the experiment was defined as the period from 2 days before feeding to 26 days after birch introduction. But *T_1/2birch_* was estimated based on the linear model estimated only for the period Day 1 to Day 10 after birch feeding, while *T_1/2lichen_* was estimated only for the period Day 13 to Day 24. The parameters of the linear model were estimated using an iteratively reweighted least squares (IRLS) algorithm implemented in the function *rlm* of the R package MASS (Venables & Rippley 2002).

## 3 Results

### Reindeer diet composition

Seven seed plant families representing more than one percent of the average reindeer diet were detected with both markers (Figure 2). Including rare families, 16 families were detected in total by *Sper01* and only 9 by *Euka02*. There was not a perfect match between diet composition retrieved from faeces and the list of plant taxa composing the pelleted feed as indicated by the manufacturer (Table S1; Figure 2A, B). Using the *Sper01* primers, four out of the six plant families composing the pelleted feed could be identified in reindeer faeces, namely Poaceae, Brassicaceae, Fabaceae and Pinaceae. Amplifications carried out with the *Euka02* primers allowed to detect and identify three of the six plant families (Poaceae, Brassicaceae and Fabaceae).

**Figure 2.**
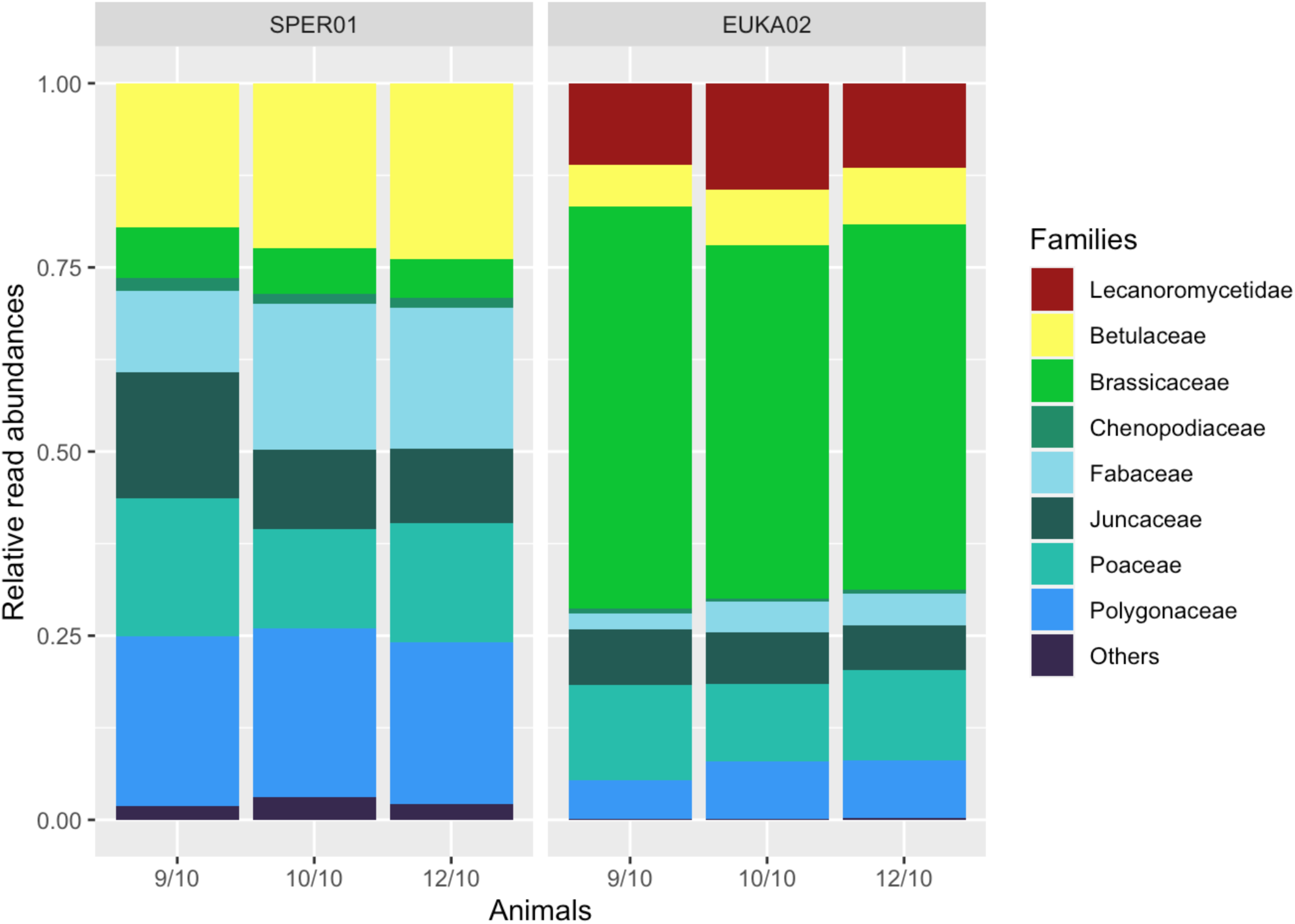
Relative read abundance (RRA) of plant families detected with the *Sper01* primers (left), and plant families and lichen (subclass Lecanoromycetidae) detected with the *Euka02* primer set (right) from faeces of three reindeer (9/10, 10/10 and 12/10).

The three animals consumed all the 50 grams of birch twigs offered at the start of the experiment, as well as all the amounts of lichen biomass offered at subsequent feedings, well within the time frame of 2 hours. Birch was identified as all MOTUs belonging the Betulaceae family, while lichen as all MOTUs belonging to the subclass Lecanoromycetidae. With the *Sper01* primers, a single MOTU belonging to the Betulaceae family was detected and Betulaceae was the most abundant plant family, representing 22% of the average reindeer diet. With the *Euka02* primers, Betulaceae represented 7% of the average diet with two MOTUs detected, which were collapsed before estimating birch DNA half-life detectability. Betulaceae DNA represented at maximum 90% of the reindeer diet during the experiment with *Sper01*, and 53% with *Euka02*. The subclass Lecanoromycetidae, only detectable with the *Euka02* primers, represented 12% of the average diet and at maximum 70% after the last lichen meal. Two MOTUs were identified as Lecanoromycetidae, also collapsed prior to estimating lichen DNA half-life detectability.

### Quantitative performance

The linear relationship between the amount of ingested food biomass and RRA in reindeer faeces was estimated for lichen, for which an increasing amount of biomass was fed during the experiment (20, 500, and 2,000 grams, Figure 3). The slopes for the three reindeer were all positive and significantly different from zero (t-test, p-value < 4.10^-5^). The intercept of the model was not different from zero (t-test, p-value = 0.126), which was expected given that the absence of food should result in the absence of DNA detection. The model had an adjusted coefficient of determination R^2^_adj_ of 0.98, and the residuals of the model were not significantly deviating from the normal distribution (Shapiro-Wilk test, p-value = 0.057).

**Figure 3.**
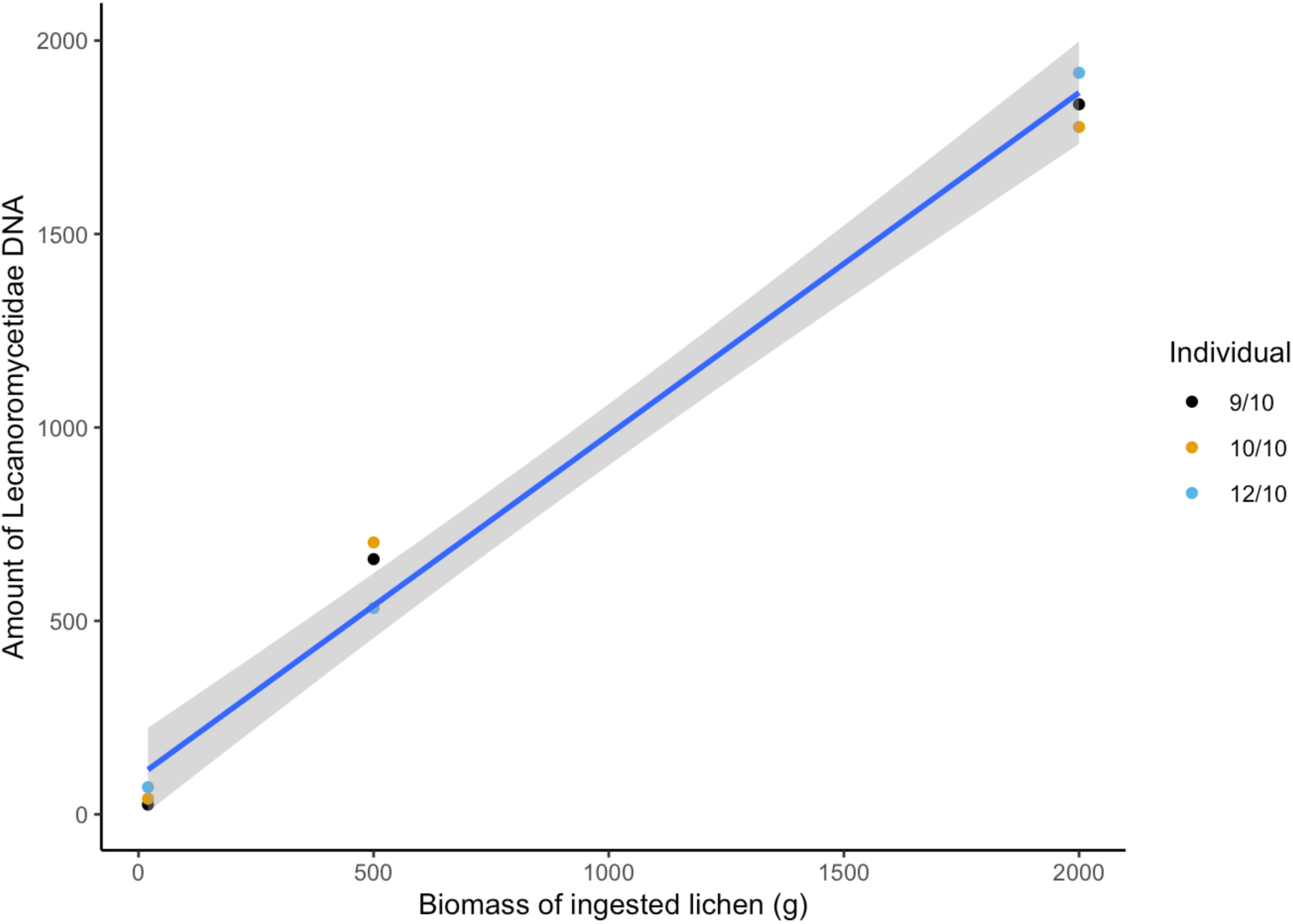
Relationship between the amount of lichen DNA (subclass Lecanoromycetidae), detected with the *Euka02* primers in the reindeer faeces, and ingested lichen biomass (linear regression model, R^2^ = 0.90). The amount of lichen DNA corresponds to the Lecanoromycetidae read abundance after accounting for the average relative read abundance of Brassicaceae in the *Euka02* dataset as well as the total amount of pelleted feed eaten by the reindeer.

### DNA half-life detectability

Average DNA half-life detectability for birch was estimated to 30.2 hours (95%CI: 22.2– 32.2) with the *Sper01* primers (R^2^ = 0.68, Table 1), and 30.4 hours (95%CI: 27.6–33.2) with the *Euka02* primer set (R^2^ = 0.70, Table 1). In line with our expectations, DNA amount of the more digestible lichen in faeces decayed faster compared to birch – 14.1 hours on average (95%CI: 13.0–15.2, R^2^ = 0.92, Table 1). T_1/2_ was not distinguishable among the three animals according to the 95% confidence intervals. T_1/2_ estimated for birch was also not distinguishable between the two primer sets. On average, lichen DNA disappeared from the faeces twice as quickly compared to birch. However, both birch and lichen DNA from the experimental feeding remained detectable over at least 26 days (i.e. until animals were released into their outdoors pasture enclosure). Birch (with both primer sets) and lichen were also both detected in reindeer faeces collected before or right at the start of the experiment, albeit in much smaller proportions (Figure 4A; S3). Moreover, a ∼10% (*Sper01*) and ∼1% (*Euka02*), respectively, increase of the DNA amount of birch was observed at 11–13 days post-feeding, although variability in DNA detectability was high (Figure 4).

**Figure 4.**
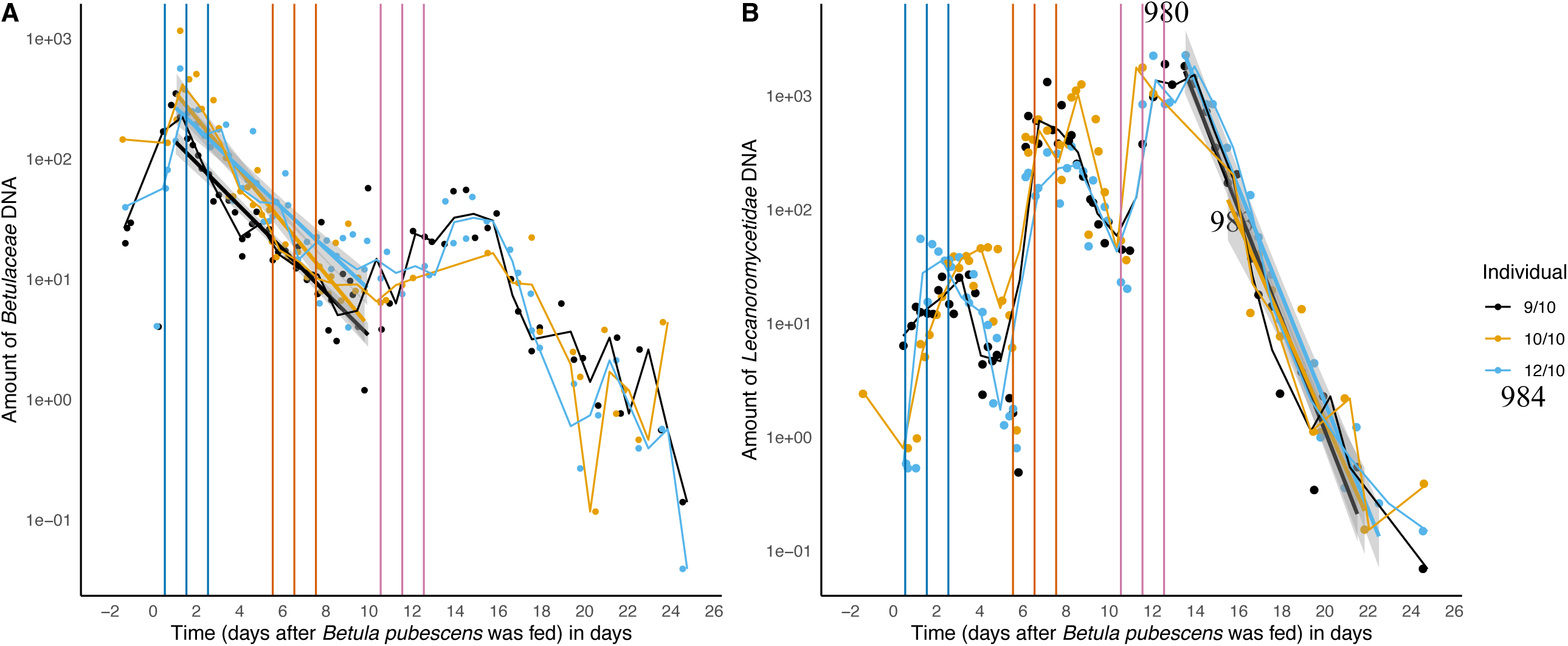
Dynamics of birch (*Betula pubescens*) and lichen (mainly *Cladonia stellaris*) DNA in faecal samples from the three individual female Eurasian tundra reindeer (*Rangifer tarandus tarandus*), detected with the *Euka02* primer sets for birch (A) and lichen (B). Solid lines represent the linear regression used to estimate DNA half-life detectability for each individual reindeer. Time 0 corresponds to the time birch was fed to the reindeer (samples collected prior to time 0 are the reference faecal samples before the experiment). Triple vertical lines in colour correspond to the progressive introduction of lichen in the reindeer diet (20 g, blue lines; 500 g, orange lines; 2,000 g, pink lines).

**Table 1.** Birch and lichen DNA half-life detectability in reindeer faeces, based on the linear model and the log-transformed sequence abundance data. DNA half-life detectability for birch was estimated with two primer sets: seed plants (*Sper01*) and eukaryotes (*Euka02*). CI indicates the 95% confidence intervals.

| Primer set | Birch DNA |  | Birch DNA |  | Lichen DNA |  |
| --- | --- | --- | --- | --- | --- | --- |
|  | *Sperm01* |  | *Euka02* |  | *Euka02* |  |
| Animal | Mean | CI (95%) | Mean | CI (95%) | Mean | CI (95%) |
| 9/10 | 32.0 h | 28.4 - 35.7 | 31.7 h | 26.7 - 36.8 | 13.2 h | 11.3 - 15.00 |
| 10/10 | 31.3 h | 27.5 - 35.2 | 27.5 h | 23.1 - 31.9 | 14.3 h | 11.0 - 17.6 |
| 12/10 | 27.4 h | 24.0 - 30.8 | 32.2 h | 27.0 - 37.5 | 14.7 h | 12.9 - 16.3 |
| All | 30.2 h | 28.2 - 32.2 | 30.4 h | 27.6 - 33.2 | 14.1 h | 13.0 - 15.2 |

## 4 Discussion

In line with our expectations, DNA metabarcoding was found to provide a reliable detection of the known plant and lichen components of the reindeer diet in our experimental set-up. Using both plant-specific and eukaryote-wide primer sets, we were able to detect food quantities of only 50 gr of birch and 20 gr of lichen, representing ∼2% of the daily food biomass intake, thereby confirming the high sensitivity of the method. This is supported by at least one other study showing that dietary items occurring at amounts as low as 1% of the total diet may be detected in faeces of the herbivore desert woodrat (*Neotoma lepida*) (Stapleton et al. 2022). Deagle et al. (2005) also show that small amounts of squid (6% of the total biomass intake) were consistently detectable in the faeces of carnivores Steller sea lions (*Eumetopias jubatus*). This is encouraging, but we still need a validation about whether or not there is a limit of detection sensitivity of DNA metabarcoding when applied to more diverse, natural diets.

We also detected a broader set of plant molecular operational taxonomic units (MOTUs) in the reindeer faeces, including some that were not part of the initial list of ingredients of the reindeer pelleted feed (cf Table S2). Some of these additional plant taxa were most likely consumed by reindeer prior to the experiment, when animals had access to natural pastures in the outdoor enclosure, as our own data show plant DNA remains detectable in the faeces for weeks. In contrast, two plant families, Amaranthaceae (the Amarant family) and Arecaceae (palm trees), both part of the listed ingredients of the reindeer pellets were never detected with any of the two primer sets. Animal feed is typically highly processed, therefore resulting in highly degraded DNA. This probably also explains why the fresh birch tissues represented 22% of the average plant diet with *Sper01* although only 50 gr were fed to the reindeer. Moreover, the actual composition of the feed might vary among batches, as suggested by the detection of Polygonaceae (plants of the buckwheat family) in reindeer faeces throughout the experiment. This family was not listed as a potential ingredient, yet such continuous ingestion from alternative sources by the three reindeer was highly unlikely given our experimental set up (i.e., snow-covered grounds and continuous monitoring of reindeer and cleaning of enclosures).

As initially predicted, our results show that the amount of lichen DNA in reindeer faeces positively correlated with the ingested lichen biomass, suggesting that DNA metabarcoding could enable the quantitative assessment of dietary proportions in LMH. DNA metabarcoding quantitative performance has been a contentious topic (Pompanon et al. 2012; Deagle et al. 2019), but also challenging to address due to the methodological difficulties of carrying out feeding trials. Currently, our best understanding comes from experiments on marine predators such as penguins and pinnipeds fed relatively simple diets and showing a mismatch between ingested biomass and DNA dietary proportions in faeces (Deagle et al. 2005; Deagle & Tollit 2007; Deagle et al. 2010). The few recent feeding trials focusing on herbivores report either good (Willerslev et al. 2014), or poor (Neby et al. 2021; Stapleton et al. 2022) agreement between sequence read abundances and ingested biomass. On the other hand, studies report good agreement in dietary proportions, when comparing DNA metabarcoding to the results from other methods (i.e. microhistology, Soininen et al. 2009, 2015; Nichols et al. 2016; stable isotope ratios, Kartzinel et al. 2015). Akin to findings from Willerslev et al. (2014), revealing significant correlation (R^2^ = 0.75) between the fraction of forbs in sheep diet and RRAs, we show that the DNA amount of lichen in faeces positively correlates with the amount of ingested biomass. This is encouraging, but it remains to be demonstrated whether this result is generalizable. We used a highly controlled set up and our sample size is limited. Moreover, we had the opportunity to use the constant proportions of plants in the baseline diet of pelleted feed to correct lichen proportions, also further corrected by the actual amount of pellets eaten by the reindeer every day. This contributed to improve the linearity of the relationship between the DNA amount and the ingested biomass of lichen. But typically, these parameters are not available for wild-ranging animals, and we still might be limited in the interpretation of dietary proportions derived from studies in natural settings. Nevertheless, DNA metabarcoding does seem to retain a quantitative signal suggesting there is value in using RRA in dietary studies (see also Deagle et al. 2019). However, one major shortcoming here is that we were only able to assess the relationship between ingested food biomass and sequence read abundance for lichen, while testing DNA metabarcoding quantitative performance typically requires a composite diet, consisting of a mixture of food items with different biomass proportions (e.g. Neby et al. 2021; Stapleton et al. 2022). But low availability of plant biomass in winter as well as the low acceptance of unfamiliar food by reindeer made it challenging to further optimise the experimental set up. Any follow-up experimentation might therefore benefit from a closer collaboration with animal feed producers or reindeer herders for the optimisation of the meal design. Meanwhile, we endorse certain confidence in using DNA metabarcoding diet data quantitatively, at least in the case of LMH, not least based on the results of the present study.

DNA from the more digestible lichen decayed significantly faster compared to birch, thus validating our second hypothesis. The amount of birch DNA detectable in faeces was reduced by half every 30 hours, compared to every 14 hours for lichen, suggesting risk for underestimating lichen proportions comparatively to the birch/shrub components of the reindeer diet. Our results need to be replicated as differences in T_1/2_ could also be influenced by additional factors such as the meal size as well as ambient temperature (Hosseini et al. 2008; Thalinger et al. 2017; Uiterwaal & DeLong 2020), the effects of which were not evaluated here. It is important to assess to what extent DNA half-lives of essential dietary items in ruminants are predictable, so that alleviation measures such as the application of correction factors could be used (Greenstone et al. 2010; Gagnon et al. 2011; Uiterwaal & DeLong 2020). No differences were observed in birch DNA half-life detectability between the two primer sets – 30.4 hours with *Euka02* versus 30.2 hours with *Sper01* (see Table 1), despite the fact that the *Euka02* DNA fragment size for birch was 102 bp long compared to only 61 bp for the *Sper01* fragment. These results imply that targeting fragments within the range 50– 100 bp should provide comparable time window estimates, if the food items have similar digestibility. Overall, DNA detectability half-life values for birch and lichen were within the lower range of the expected time window of 24–48 hours with values comparable to estimates in mammals with supposedly faster digestion – e.g. rats and hedgehogs predating on frogs (T_1/2_ ∼21 hours), detected with a DNA fragment size of ∼130 bp (Egeter et al. 2014). However, whether the same processes drive DNA decay dynamics remains unclear. The digestive systems of carnivores and herbivores are fundamentally different, due to differences in the digestibilities of their diet and the anatomy of their gastrointestinal tract affecting gut fill and transit times (De Cuyper et al. 2020). Thus, building a general understanding of the mechanisms driving dietary DNA detectability would require more extensive experimentation across a wider range of mammalian species differing in size, digestive systems, diet breadth and quality.

But our findings raise the question about the interpretation with DNA methods of the temporal dynamics of herbivorous diets. In general, dietary DNA started to appear in faeces in less than 24 hours post-feeding, independently of the primer set, thus corroborating the idea that DNA metabarcoding detects very recent feeding events (Deagle et al. 2005; Franz et al. 2023). Yet, our data also show that trace amounts of birch DNA were detectable over the duration of our experiment (i.e. almost four weeks). This suggests that a single reindeer scat represents both a temporal snapshot of the most recent diet and traces of meals ingested over a period of three to four weeks. It also implies that assessing diet selectivity with DNA metabarcoding in highly mobile grazers, such as reindeer, might be challenging given the large spatial scale and habitat heterogeneity potentially covered by the animals in such a time period. The likely explanation for the long DNA detectability lies in the complexity of the digestive process in ruminants, involving rumination, rumen fill and stratification of digesta, selective retention of food particles based on their size as well as differential passages of fluid versus solid fractions of the diet (Claus et al. 2010; Lechner et al. 2010; Lauper et al. 2013). Typically, the highly digestible portions of the food are retained in the rumen for fermentation for longer periods, while indigestible fractions, such as lignin, are rapidly cleared through the digestive tract (Moyo & Nsahlai 2017), likely explaining the rapid detection of dietary DNA few hours after ingestion, as well as its continuous detection over several weeks. Many ruminants also have a stratified rumen content, where the gas, liquid, and solid components of digesta are structured into distinct layers according to density (Tschuor & Clauss 2008), leading to food particles entrapment into the floating fibermat, potentially delaying their passage (Moyo & Nsahlai 2017). However, as in our case pelleted feed represented reindeer’s bulk diet, it is unlikely that such homogenous food can result in stratification of the rumen content. But fractions of the digesta could also be retained for further fermentation within the caecum (Sørmo et al. 1997). Thus, the prolonged DNA detection as well as the small peak in birch DNA (∼1% with *Euka02*, ∼10% with *Sper01*, Figure 3A, S3) seen approximately two weeks after the start of the experiment, may be due to a sequential release of digesta from the same meal. The relatively long dietary DNA detection here also suggests that the sampling window of DNA metabarcoding is comparable to the sampling window of other methods. For instance, carbon and nitrogen stable isotope ratios turnover in reindeer blood serum is typically 1 to 2 weeks, and 3 to 4 weeks for the blood clot (Hiltunen et al. 2022). Yet, the two methods remain highly complementary, since DNA metabarcoding estimates diet composition and diversity, while stable isotope ratios indicate the degree of trophic enrichment, or the part of the diet that is actually assimilated into the host’s tissues. On the other hand, it is possible with DNA metabarcoding to restrict diet analysis to the most recent feeding events, for example by increasing the size of the target DNA fragment, (e.g. Kamenova et al. 2018), which for plants could correspond to amplifying the whole length of the *trn*L intron (cf Taberlet et al. 2007).

## 5 Conclusion

DNA metabarcoding has revolutionised our capacity to generate high-resolution data on diet composition. However, we still lack general understanding about its methodological limitations. In three experimentally fed reindeer, DNA metabarcoding enabled the detection of low-abundant, recently ingested and highly digested plants and lichen. Our results also show that the proportion of ingested lichen biomass positively correlates with the amount of DNA, suggesting potential for determining dietary proportions from DNA metabarcoding sequence data in herbivores. Most importantly, dietary DNA remained detectable in faeces for three to four weeks post-feeding, thus providing a far more integrative overview of a ruminant’s diet than previously expected. This result might question our current understanding of the digestive process in ruminants. We call for more replication studies to advance knowledge about the method and improve the interpretation of DNA metabarcoding diet data.

## Acknowledgements

We thank Tove Aagnes Utsi for the stimulating discussions that led to the development of this project. We also thank Hans Lian, Hans Arne Solvang, and Renate Thorvaldsen for all the technical help and advice provided throughout the experiment, and Gabriela Wagner for her invaluable input on reindeer physiology and behaviour. Many thanks to Nora Slåttebrekk and Jaymee van Dalum for helping with running the experiment. This project received funding from the Research Council of Norway through project number 257642 REININ “Reindeer interactions from plants and birds to humans: balancing the odds of climate change” (PI: Galina Gussarova). Stefaniya Kamenova was also supported by European Commission Research and Innovation Action Number 869471 (CHARTER “Drivers and Feedbacks of Changes in Arctic Terrestrial Biodiversity”, PI: Bruce Forbes) and the Research Council of Norway project 315454 (PRISM “Understanding climate change impacts in an Arctic ecosystem: an integrated approach through the prism of Svalbard reindeer”, PI: Leif Egil Loe).

## Author contributions

All authors contributed to ideas and design of the study. MAS and LPF provided expertise and access to reindeer and animal facilities at UiT. PM and SK carried out the animal experiment and sampling, together with LR. PM and SK carried out DNA metabarcoding analyses and bioinformatics. ÉC analysed the data with input from PM, and SK wrote the initial draft. All authors contributed critically to developing and finalising the manuscript.

## Permits

Handling of animals in this experiment did not require any authority permit, as defined by the conditions listed in §2, part 5, items ‘a-f’ of the “Regulations on the use of animals in experiments” (“Forskrift om bruk av dyr i forsøk”, FOR-2015-06-18-761 https://lovdata.no/dokument/SF/forskrift/2015-06-18-761), as regulated by the Norwegian Food Safety Authority and conforming with regulations in EU Directive 2010/63/EU (https://eurlex.europa.eu/LexUriServ/LexUriServ.do?uri=OJ:L:2010:276:0033:0079:en:PDF), Chapter 1, Article 1, item 5 a-f.

## Data availability statement

Demultiplexed sequence datasets as well as datasets used for statistical analyses, metadata and scripts are available at https://github.com/RENIN-Project/Feeding-Experiment/tree/main. Additional information about the data used for statistical analyses are provided in the Supplemental material. Raw sequence files as well as bioinformatic scripts will be made available upon publication.

## References

Ando H, Mukai H, Komura T, Dewi T, Ando M, Isagi Y (2020) Methodological trends and perspectives of animal dietary studies by noninvasive fecal DNA metabarcoding. Environmental DNA 2, 391–406. DOI: 10.1002/edn3.117

Barboza PS, Peltier TC, Forster RJ (2006) Ruminal fermentation and fill change with season in an arctic grazer: responses to hyperphagia and hypophagia in muskoxen (*Ovibos moschatus*). Physiological and Biochemical Zoology 79, 497–513. DOI: 10.1086/501058

Bendich AJ (1987) Why do chloroplasts and mitochondria contain so many copies of their genome? Bioessays 6, 279–82. DOI: 10.1002/bies.950060608.

Bennett MD, Leitch IJ (2012) Plant DNA C-values Database. Available at: http://data.kew.org/cvalues/

Boyer F, Mercier C, Bonin A, Le Bras Y, Taberlet P, Coissac E (2016) *obitools*: a unix-inspired software package for DNA metabarcoding. Molecular Ecology Resources 16, 176–82. DOI: 10.1111/1755-0998.12428

Chen Y, Giles KL, Payton ME, Greenstone MH (2000) Identifying key cereal aphid predators by molecular gut analysis. Molecular Ecology 9, 1887–1898. DOI: 10.1046/j.1365-294x.2000.01100.x.

Clauss M, Hume I, Hummel J (2010) Evolutionary adaptations of ruminants and their potential relevance for modern production systems. Animal 4, 979–992. DOI: 10.1017/S1751731110000388

Crater AR, Barboza PS, Forster RJ (2007) Regulation of rumen fermentation during seasonal fluctuations in food intake of muskoxen. Comparative Biochemistry and Physiology Part A: Molecular and Integrative Physiology 146, 233–241. DOI: 10.1016/j.cbpa.2006.10.019

Deagle BE, Chiaradia A, McInnes J, Jarman SN (2010) Pyrosequencing faecal DNA to determine diet of little penguins: is what goes in what comes out? Conservation Genetics 11, 2039–2048. DOI: 10.1007/s10592-010-0096-6

Deagle BE, Eveson JP, Jarman SN (2006) Quantification of damage in DNA recovered from highly degraded samples – a case study on DNA in faeces. Frontiers in Zoology 3, 11. DOI: 10.1186/1742-9994-3-11

Deagle BE, Thomas AC, McInnes JC, Clarke LJ, Vesterinen EJ, Clare EL, Kartzinel TR, Eveson JP (2019) Counting with DNA in metabarcoding studies: How should we convert sequence reads to dietary data? Molecular Ecology 28, 391–406. DOI: 10.1111/mec.14734

Deagle BE, Thomas AC, Shaffer AK, Trites AW, Jarman SN (2013) Quantifying sequence proportions in a DNA-based diet study using Ion Torrent amplicon sequencing: which counts count? Molecular Ecology Resources 13, 620–633. DOI: 10.1111/1755-0998.12103

Deagle BE, Tollit DJ (2007) Quantitative analysis of prey DNA in pinniped faeces: potential to estimate diet composition? Conservation Genetics 8, 743–747. DOI: 10.1007/s10592-006-9197-7

Deagle BE, Tollit DJ, Jarman SN, Hindell MA, Trites AW, Gales NJ (2005) Molecular scatology as a tool to study diet: analysis of prey DNA in scats from captive Steller sea lions. Molecular Ecology 14, 1831–1842. DOI: 10.1111/j.1365-294X.2005.02531.x

Dearing MD, Kohl KD (2017) Beyond fermentation: other important services provided to endothermic herbivores by their gut microbiota. Integrative and Comparative Biology 57, 723– 731. DOI: 10.1093/icb/icx020

De Cuyper A, Meloro C, Abraham AJ, Müller DWH, Codron D, Janssens GPJ, Clauss M (2020) The uneven weight distribution between predators and prey: Comparing gut fill between terrestrial herbivores and carnivores. Comparative Biochemistry and Physiology Part A: Molecular & Integrative Physiology 243, 110683. DOI: 10.1016/j.cbpa.2020.110683

Dehority BA (2003) Rumen microbiology. Nottingham University Press, Nottingham.

Egeter B, Bishop PJ, Robertson BC (2014) Detecting frogs as prey in the diets of introduced mammals: a comparison between morphological and DNA-based diet analyses. Molecular Ecology Resources 15, 306–316. DOI: 10.1111/1755-0998.12309

Franz M, Whyte L, Atwood TC, Menning D, Sonsthagen SA, Talbot SL, Laidre KL, Gonzalez E, McKinney MA (2023) Fecal DNA metabarcoding shows credible short-term prey detections and explains variation in the gut microbiome of two polar bear subpopulations. Marine Ecology Progress Series 704, 131–147. DOI: 10.3354/meps14228

Gagnon A-È, Doyon J, Heimpel GE, Brodeur J (2011) Prey DNA detection success following digestion by intraguild predators: influence of prey and predator species. Molecular Ecology Resources 11, 1022–1032. DOI: 10.1111/j.1755-0998.2011.03047.x

Garnick S, Barboza PS, Walker JW (2018) Assessment of animal-based methods used for estimating and monitoring rangeland herbivore diet composition. Rangeland Ecology and Management 71, 449–457. DOI: 10.1016/j.rama.2018.03.003

Greenstone MH, Payton ME, Weber DC, Simmons AM (2014) The detectability half-life in arthropod predator–prey research: what it is, why we need it, how to measure it, and how to use it. Molecular Ecology 23, 3799–3813. DOI: 10.1111/mec.12552

Greenstone MH, Rowley DL, Weber DC, Payton ME, Hawthorne DJ (2007) Feeding mode and prey detectability half-lives in molecular gut-content analysis: an example with two predators of the Colorado potato beetle. Bulletin of Entomological Research 97, 201–209. DOI: 10.1017/S000748530700497X

Greenstone MH, Szendrei Z, Payton ME, Rowley DL, Coudron TC, Weber DC (2010) Choosing natural enemies for conservation biological control: use of the prey detectability half-life to rank key predators of Colorado potato beetle. Entomologia Experimentalis et Applicata 136, 97–107. DOI: 10.1111/j.1570-7458.2010.01006.x

Guardiola M, Uriz MJ, Taberlet P, Coissac E, Wangensteen OS, Turon X (2015) Deep-Sea, deep-sequencing: metabarcoding extracellular DNA from sediments of marine canyons. PLoS ONE 10(10): e0139633. DOI: 10.1371/journal.pone.0139633

Hiltunen TA, Stien A, Väisänen M, Ropstad E, Aspi JO, Welker JM (2022) Svalbard reindeer winter diets: Long-term dietary shifts to graminoids in response to a changing climate. Global Change Biology 28, 7009–7022. DOI: 10.1111/gcb.16420

Hofmann RR (1989) Evolutionary steps of ecophysiological adaptation and diversification of ruminants: a comparative view of their digestive system. Oecologia 78, 443–357.

Hofmann RR (2011) Functional and comparative digestive system anatomy of Arctic ungulates. Rangifer 20, 71. DOI: 10.7557/2.20.2-3.1504

Hosseini R, Schmidt O, Keller MA (2008) Factors affecting detectability of prey DNA in the gut contents of invertebrate predators: A polymerase chain reaction-based method. Entomologia Experimentalis et Applicata 126, 194–202. DOI: 10.1111/j.1570-7458.2007.00657.x

Hungate RE (1975) The rumen microbial ecosystem. Annual Review of Ecology and Systematics 6.1, 39–66.

Ingólfsdóttir K (2002) Molecules of interest. Usnic acid. Phytochemistry 61, 729–736.

Jernsletten J-L, Klokov K (2002) Sustainable reindeer husbandry. Arctic Council 2000–2002. University of Tromsø, Centre for Sami Studies, Tromsø, p. 157.

Jorns T, Craine J, Towne EG, Knox M (2020) Climate structures bison dietary quality and composition at the continental scale. Environmental DNA 2, 77–90. DOI: 10.1002/edn3.47

Kamenova S, Mayer R, Rubbmark OR, Coissac É, Plantegenest M, Traugott M (2018) Comparing three types of dietary samples for prey DNA decay in an insect generalist predator. Molecular Ecology Resources 18, 966–973. DOI: 10.1111/1755-0998.12775

Kartzinel TR, Chen PA, Coverdale TC, Erickson DL, Kress WJ, Kuzmina ML, Rubenstein DI, Wang W, Pringle RM (2015) DNA metabarcoding illuminates dietary niche partitioning by African large herbivores. Proceedings of the National Academy of Sciences of the United States of America 112, 8019– 8024. DOI: 10.1073/pnas.1503283112

Kay RNB, von Engelhardt W, White RG (1980) The digestive physiology of wild ruminants. Ruckebush Y, Thivend P (Eds.) Digestive physiology and metabolism in ruminants, MTP Press, Lancaster, UK, pp. 743–761.

Lauper M, Lechner I, Barboza PS, Collins WB, Hummel J, Codron D, Clauss M (2013) Rumination of different-sized particles in muskoxen (*Ovibos moschatus*) and moose (*Alces alces*) on grass and browse diets, and implications for rumination in different ruminant feeding types. Mammalian Biology 78, 142–152. DOI: 10.1016/j.mambio.2012.06.001

Lechner I, Barboza P, Collins W, Fritz J, Günther D, Hattendorf B, Hummel J, Südekum K-H, Clauss M (2010) Differential passage of fluids and different-sized particles in fistulated oxen (*Bos primigenius* f. *taurus*), muskoxen (*Ovibos moschatus*), reindeer (*Rangifer tarandus*) and moose (*Alces alces*): Rumen particle size discrimination is independent from contents stratification. Comparative Biochemistry and Physiology Part A: Molecular and Integrative Physiology 155, 211–222. DOI: 10.1016/j.cbpa.2009.10.040

Lin Z et al. (2019) Biological adaptations in the Arctic cervid, the reindeer (*Rangifer tarandus*). Science 364, eaav6312. DOI: 10.1126/science.aav6312

Mathiesen SD, Aagnes Utsi TH, Sørmo W (1999) Forage chemistry and the digestive system in reindeer (*Rangifer tarandus tarandus*) in northern Norway and on South Georgia. Rangifer 19, 91–101. DOI: 10.7557/2.19.2.285

Mathiesen SD, Haga Ø, Kaino T, Tyler N (2000a) Diet composition, rumen papillation and maintenance of carcass mass in female Norwegian reindeer (*Rangifer tarandus tarandus*) in winter. Journal of Zoology 251, 129–138. DOI: 10.1111/j.1469-7998.2000.tb00598.x

Mathiesen SD, Sørmo W, Haga ØE, Norberg HJ, Utsi THA, Tyler NJC (2000b) The oral anatomy of Arctic ruminants: coping with seasonal changes. Journal of Zoology 251, 119–128. DOI: 10.1111/j.1469-7998.2000.tb00597.x

Mitchell G, Wilson PJ, Manseau M, Redquest B, Patterson BR, Rutledge LY (2022) DNA metabarcoding of faecal pellets reveals high consumption of yew (*Taxus* spp.) by caribou (Rangifer tarandus) in a lichen-poor environment. FACETS 7, 701–717. DOI: 10.1139/facets-2021-0071

Meyer, P (2019) Validation of DNA metabarcoding as a tool for diet analysis in reindeer (*Rangifer tarandus tarandus* L.) MSc Thesis, University of Oslo. http://urn.nb.no/URN:NBN:no-73746

Moyo M, Gueguim Kana EB, Nsahlai IV (2017) Modelling of digesta passage rates in grazing and browsing domestic and wild ruminant herbivores. South African Journal of Animal Science 47. DOI: 10.4314/sajas.v47i3.13

Moyo M, Nsahlai IV (2017) Rate of passage of digestain ruminants; are goats different? In: Goat science. Kukovics S (ed.). pp: 39–74. IntechOpen, London. DOI: 10.5772/intechopen.69745

Müller-Wille L, Heinrich D, Lehtola V-P, Aikio P, Konstantinov Y, Vladimirova V (2006) Dynamics in human–reindeer relations: reflections on prehistoric, historic and contemporary practices in northernmost Europe. In Forbes BC, Bölter M, Müller-Wille L, Hukkinen J, Müller F, Gunslay N, Konstantinov Y (Eds.), Reindeer management in northernmost Europe: linking practical and scientific knowledge in social-ecological systems. Ecological Studies 184, 11–25. Berlin: Springer-Verlag.

Neby M, Kamenova S, Devineau O, Ims RA, Soininen EM (2021) Issues of under-representation in quantitative DNA metabarcoding weaken the inference about diet of the tundra vole *Microtus oeconomus*. PeerJ 9, e11936. DOI: 10.7717/peerj.11936

Nichols RV, Åkesson M, Kjellander P (2016) Diet assessment based on rumen contents: a comparison between DNA metabarcoding and macroscopy. PLoS ONE 11(6): e0157977. DOI: 10.1371/journal.pone.0157977

Palo RT (1985) Chemical defense in birch: Inhibition of digestibility in ruminants by phenolic extracts. Oecologia 68, 10–14. DOI: 10.1007/BF00379465

Palo RT (1993) Usnic acid, a secondary metabolite of lichens and its effect on in vitro digestibility in reindeer. Rangifer 13, 39–43. DOI: 10.7557/2.13.1.1071

Pansu J, Gayton JA, Potter AB, Atkins JL, Daskin JH, Wursten B, Kartzinel TR, Pringle RM (2019) Trophic ecosystem of large herbivores in a reassembling African ecosystem. Journal of Ecology 107, 1355–1376. DOI: 10.1111/1365-2745.13113

Paula DP, Linard B, Andow DA, Sujii ER, Pires CSS, Vogler AP (2014) Detection and decay rates of prey and prey symbionts in the gut of a predator through metagenomics. Molecular Ecology Resources 15, 880–892. DOI: 10.1111/1755-0998.12364

Picard M, Papaïx J, Gosselin F, Picot D, Bideau E, Baltzinger C (2015) Temporal dynamics of seed excretion by wild ungulates: implications for plant dispersal. Ecology and Evolution 5, 2621–2632. DOI: 10.1002/ece3.1512

Pompanon F, Deagle BE, Symondson WOC, Brown DS, Jarman SN, Taberlet P (2012) Who is eating what: diet assessment using next generation sequencing. Molecular Ecology 21, 1931–1950. DOI: 10.1111/j.1365-294X.2011.05403.x

Pope PB, Mackenzie AK, Gregor I, Smith W, Sundset MA, McHardy AC, et al. (2012) Metagenomics of the Svalbard reindeer rumen microbiome reveals abundance of polysaccharide utilization loci. PLoS ONE 7(6): e38571. DOI: 10.1371/journal.pone.0038571

Royston P (1982) Algorithm AS 181: The W test for Normality. Applied Statistics 31, 176–180. DOI:10.2307/2347986.

Salgado-Flores A, Hagen LH, Ishaq SL, Zamanzadeh M, Wright ADG, Pope PB, Sundset MA (2016) Rumen and cecum microbiomes in reindeer (*Rangifer tarandus tarandus*) are changed in response to a lichen diet and may affect enteric methane emissions. PLoS ONE 11(5): e0155213. DOI: 10.1371/journal.pone.0155213

Scasta JD, Jornsa T, Derner JD, Lake S, Augustine DJ, Windh JL, Smith TL (2019) Validation of DNA metabarcoding of fecal samples using cattle fed known rations. Animal Feed Science and Technology 255, 114219. DOI: 10.1016/j.anifeedsci.2019.114219

Soininen EM, Gauthier G, Bilodeau F, Berteaux D, Gielly L, Taberlet P, et al. (2015) Highly overlapping winter diet in two sympatric lemming species revealed by DNA metabarcoding. PLoS ONE 10(1): e0115335. DOI: 10.1371/journal.pone.0115335

Soininen EM, Valentini A, Coissac E, Miquel C, Gielly L, Brochmann C, Brysting AK, Sønstebø JH, Ims RA, Yoccoz NG, Taberlet P (2009) Analysing diet of small herbivores: the efficiency of DNA barcoding coupled with high-throughput pyrosequencing for deciphering the composition of complex plant mixtures. Frontiers in Zoology 6, 16. DOI: 10.1186/1742-9994-6-16

Spitzer R, Coissac E, Felton A, Fohringer C, Juvany L, Landman M, Singh NJ, Taberlet P, Widemo F, Cromsigt JPGM (2021) Small shrubs with large importance? Smaller deer may increase the moose-forestry conflict through feeding competition over *Vaccinium* shrubs in the field layer. Forest Ecology and Management 480, 118768. DOI: 10.1016/j.foreco.2020.118768

Stapleton TE, Weinstein SB, Greenhalgh R, Dearing MD (2022) Successes and limitations of quantitative diet metabarcoding in a small, herbivorous mammal. Molecular Ecology Resources 22, 2573–2586. DOI: 10.1111/1755-0998.13643

Steuer P, Südekum K-H, Müller DWH, Franz R, Kaandorp J, Clauss M, Hummel J (2011) Is there an influence of body mass on digesta mean retention time in herbivores? A comparative study on ungulates. Comparative Biochemistry and Physiology Part A: Molecular and Integrative Physiology 160, 355–364. DOI: 10.1016/j.cbpa.2011.07.005

Storeheier PV, Mathiesen SD, Schjelderup I, Tyler NJC, Olsen MA (2002a) Utilization of nitrogen and mineral rich vascular forage plants by reindeer in winter. Journal of Agricultural Science 139. DOI: 10.1017/S0021859602002344

Storeheier PV, Mathiesen SD, Schjelderup I, Tyler NJC, Olsen MA (2002b) Nutritive value of terricolous lichens for reindeer in winter. Lichenologisk 34, 247–257. DOI: 10.1006/lich.2002.0394

Storeheier PV, Van Oort BEH, Sundset MA, Mathiesen SD (2003) Food intake of reindeer in winter. The Journal of Agricultural Science 141, 93–101. DOI: 10.1017/S002185960300337X

Sundset MA, Barboza P, Green TK, Folkow LP, Blix AS, Mathiesen SD (2010). Microbial degradation of usnic acid in the reindeer rumen. Die Naturwissenschaften 97, 273–278. DOI: 10.1007/s00114-009-0639-1

Sundset MA, Kohn A, Mathiesen SD, Praesteng KE (2008) *Eubacterium rangiferina*, a novel usnic acid-resistant bacterium from the reindeer rumen. Naturwissenschaften. 95, 741–9. DOI: 10.1007/s00114-008-0381-0.

Sundset MA, Cann IKO, Mathiesen SD, Præsteng KE, Mackie RI (2007) Novel rumen bacterial diversity in two geographically separated sub-species of reindeer. Microbial Ecology 54, 424–438. DOI: 10.1007/s00248-007-9254-x

Sundset MA, Edwards JE, Cheng YF, Sensosiain RS, Fraile MN, Northwood KS, Præsteng KE, Glad T, Mathiesen SD, Wright ADG (2009) Molecular diversity of the rumen microbiome of Norwegian reindeer on natural pasture. Microbial Ecology 57, 335–348. DOI: 10.1007/s00248-008-9414-7

Sundset MA, Salgado-Flores A, Wright ADG, Pope PB (2013) The Reindeer Rumen Microbiome. In: Nelson, K. (eds) Encyclopedia of Metagenomics. Springer, New York, NY. DOI: 10.1007/978-1-4614-6418-1_664-1

Sunnerheim K, Palo RT, Theander O, Knutsson P-G (1988) Chemical defense in birch. Platyphylloside: A phenol from *Betula pendula* inhibiting digestibility. Journal of Chemical Ecology 14, 549–60. DOI: 10.1007/BF01013906

Sørmo W, Haga ØE, White R, Mathiesen SD (1997) Comparative aspects of volatile fatty acid production in the rumen and distal fermentation chamber in Svalbard reindeer. Rangifer 17, 81–95. DOI: 10.7557/2.17.2.1355

Taberlet P, Coissac E, Pompanon F, Gielly L, Miquel C, Valentini A, Vermat T, Corthier G, Brochmann C, Willerslev E (2007) Power and limitations of the chloroplast *trn*L (UAA) intron for plant DNA barcoding. Nucleic Acids Research 35, e14. DOI: 10.1093/nar/gkl938

Thalinger B, Oehm J, Obwexer A, Traugott M (2017) The influence of meal size on prey DNA detectability in piscivorous birds. Molecular Ecology Resources 17, e174–e186. DOI: 10.1111/1755-0998.12706

Tschuor A, Clauss M (2008) Investigations on the stratification of forestomach contents in ruminants: an ultrasonographic approach. European Journal of Wildlife Research 54, 627–633. DOI: 10.1007/s10344-008-0188-5

Turi JM (2002) The world reindeer livelihood – current situation, threats and possibilities. In: Kankaanpää S, Müller-Wille L, Susiluoto P, Sutinen M-L (Eds.) Northern Timberline Forests: Environmental and Socio-Economic Issues and Concerns, pp. 70–75. The Finnish Forest Research Institute, Kolari.

Uiterwaal SF, DeLong JP (2020) Using patterns in prey DNA digestion rates to quantify predator diets. Molecular Ecology Resources 20, 1723–1732. DOI: 10.1111/1755-0998.13231

van Oort B, Tyler N, Gerkema M, Folkow L, Blix AS, Stokkan K-A (2005) Circadian organization in reindeer. Nature 438, 1095–1096. DOI: 10.1038/4381095a

Venables WN, Ripley BD (2002) Modern Applied Statistics with S. Fourth Edition. Springer, New York. ISBN 0–387-95457-0

von Berg K, Traugott M, Symondson WOC, Scheu S (2008) The effects of temperature on detection of prey DNA in two species of carabid beetle. Bulletin of Entomological Research 98, 263–269. DOI: 10.1017/S0007485308006020

Weber DC, Lundgren JG (2009) Detection of predation using qPCR: Effect of prey quantity, elapsed time, chaser diet, and sample preservation on detectable quantity of prey DNA. Journal of Insect Science 9, 41. DOI: 10.1673/031.009.4101

Willerslev E, Davison J, Moora M. et al. (2014) Fifty thousand years of Arctic vegetation and megafaunal diet. Nature 506, 47–51. DOI: 10.1038/nature12921

Zinger L, Taberlet P, Schimann H, Bonin A, Boyer F, De Barba M, Gaucher P, Gielly L, Giguet-Covex C, Iribar A et al. (2019) Body size determines soil community assembly in a tropical forest. Molecular Ecolology 28, 528–543. DOI: 10.1111/mec.14919

Zoschke R, Liere K, Börner T (2007) From seedling to mature plant: *Arabidopsis* plastidial genome copy number, RNA accumulation and transcription are differentially regulated during leaf development. The Plant Journal 50, 710–722. DOI: 10.1111/j.1365-313X.2007.03084.x

