## Supplementary material for "DNA metabarcoding diet analysis in ruminants is quantitative and integrates feeding over several weeks": Table S1

*Equal contribution

### Table of Contents:

| | |
|---|---|
| **Appendix 1:** DNA metabarcoding analysis | Page 4 |
| **Appendix 2:** Bioinformatic processing of sequence data | Page 5 |
| **Appendix 3:** Data curation | Page 5 |
| **Supplemental Table S1:** Main plant ingredients of the RF-80 pelleted feed provided *ad libitum* to the reindeer | Page 7 |
| **Supplemental Table S2:** Quantity of RF-80 feed consumed every day by the three reindeer | Page 8 |
| **Supplemental Table S3:** Number of reindeer faeces collected immediately before, during and after the feeding experiment | Page 9 |
| **Supplemental Table S4:** Details about the 234 faecal time-points selected for DNA metabarcoding diet analysis | Page 11 |
| **Supplemental Table S5:** Details on the two DNA metabarcoding primer sets used for the analysis of reindeer diets | Page 17 |
| **Supplemental Table S6:** Summary of the six synthetic DNA standards used as a mock community PCR positive control with the Sper01 plant primers | Page 18 |
| **Supplemental Figure S1:** Overview of the seed plant families as detected from the reindeer faeces with the *Sper01* primer set | Page 19 |
| **Supplemental Figure S2:** Overview of the seed plant families and lichen (subclass Lecanoromycetidae) as detected from the reindeer faeces with the *Euka02* primer set | Page 20 |
| **Supplemental Figure S3:** Proportion of birch DNA (*Betula pubescence*) in | Page 21 |

| |
|---|
| reindeer faecal samples, as detected with the plant *Sper01* primer set. |

#### **Appendix 1:** DNA metabarcoding analysis

Primers were synthesized with 8 or 9 bp sequence tags (https://github.com/pheintzman/metabarcoding) at each extremity in order to assign the sequences to each sample after sequencing. PCRs were run in triplicate. Reactions were carried out in a final volume of 15 µl using the AmpliTaq Gold 360 PCR Master Mix (Thermo Fisher Scientific, USA), 0.4 µl/15 ml of bovine serum albumin (BSA, Sigma-Aldrich, USA), 0.5 µM of each primer and 2 µl of undiluted DNA. PCR conditions were as follows: Initial denaturation at 95°C for 10 min, followed by 35 cycles consisting of denaturation at 95°C for 30 s, annealing temperature of 52° (*trn*L *Sper01*) or 45° (18S *Euka02*) for 30 s, elongation at 72°C for 1 min, and a final elongation at 72°C for 7 min before a final hold at 15°C. Two PCR negative controls (ultra-pure Milli-Q water instead of DNA) per batch of 94 samples were included in order to monitor for possible contaminations. For *Sper01*, positive controls consisted of an artificially assembled mock community containing a mixture of varying proportions of six unique synthetic DNA stretches with varying GC content, homopolymers and sequence length (Table S6). For the *Euka02*, positive controls consisted in a single synthetic DNA sequence used at a concentration of 1 ng/µL (Table S6). PCR products were first pooled per primer set and purified using the QIAquick PCR Purification Kit (Qiagen, Germany). DNA concentrations from purified amplicon pools were quantified using a Qubit 2.0 fluorometer and the dsDNA HS Assay kit (Invitrogen, Life Technologies, USA), and pooled again prior to library preparation and sequencing.

#### Appendix 2: Bioinformatic processing of sequence data

Bioinformatic analyses were carried out with the OBITools program (Boyer et al. 2016)[1]. The forward and the reverse pair-end reads were aligned and merged into a consensual sequence by considering the quality of the sequence data during the alignment and the consensus computation. Only alignments with scores >50 were kept for further analyses. For each primer set, sequences were assigned to samples, and only sequences with a perfect match on tags and a maximum of two errors on primers were retained for further analyses. Primers and tags were cut off at this step. Strictly identical sequences were clustered together, while keeping the information about their distribution among samples. Sequences shorter than 10 bp and/or occurring at ≤10 reads in the whole dataset were filtered out. Taxonomic assignments were carried out using the lowest common ancestor (LCA) algorithm as implemented in the OBITools *ecotag command*. Reference database used for this annotation were constructed for each primer pair by extracting the corresponding DNA region for the relevant taxonomic groups from the European Nucleotide Archive nucleotide library using *ecoPCR* (Ficetola et al. 2010).

#### Appendix 3: Data curation

MOTUs representing less than 1% in at least one PCR replicate were discarded, thus effectively filtering out any tag jumps (tracked through the distribution and relative abundance of synthetic standards in PCR positive controls). Aberrant PCR replicates were discarded under the

---

[1]Boyer F, Mercier C, Bonin A, Le Bras Y, Taberlet P, Coissac E (2016) obitools: a unix-inspired software package for DNA metabarcoding. Molecular Ecology Resources 16, 176-82. DOI: 10.1111/1755-0998.12428; [2]Ficetola GF, Coissac E, Zundel S, Riaz T, Shehzad W, Bessière J, Taberlet P, Pompanon F (2010) An In silico Approach for the Evaluation of DNA Barcodes. BMC Genomics 11, 434. DOI: 10.1186/1471-2164-11-434.

assumption that they were the result of non-functional PCR reactions. For each sample, the Euclidean distances of its PCR replicates from their barycenters are calculated. For a given sample, if at least one of its PCR replicates has a distance greater than 0.3 (a threshold defined empirically from the observed distribution of estimated distances), the most distant replicate is discarded. This process was repeated iteratively until no more PCR replicates were removed from the data set. If only one PCR replicate per sample remained at the end, the sample was deleted from the dataset. At the end of this procedure, relative read frequencies of the remaining PCR replicates were averaged for each sample. PCR outliers could not be filtered out for 54 samples, as replicates were labelled using the same tags (Table S4). PCR filtered and non-filtered samples were nevertheless integrated into the same dataset. MOTUs matching potential contaminants were discarded (i.e. banana, 0.2% of the total reads abundance). PCR amplification success was confirmed by successfully retrieving all synthetic standard sequences.

**Table S1:** Main plant ingredients of the RF-80 pelleted feed (FK Reinfôr, Felleskjøpet, Norway) provided *ad libitum* to the reindeer during the experiment.

| Ingredient | Latin name | Family |
|---|---|---|
| Barley | *Hordeum vulgare* | Poaceae |
| Oat | *Avena sativa* | Poaceae |
| Wheat bran | *Triticum aestivum* | Poaceae |
| Rape seed | *Brassica napus* | Brassicaceae |
| Rape fibre | *Brassica napus* | Brassicaceae |
| Beet pulp | *Beta vulgaris* | Amaranthaceae |
| Soy flour | *Glycine max* | Fabaceae |
| Molasses, sugarcane | *Saccharum officinarum* | Poaceae |
| Molasses, beet | *Beta vulgaris* | Amaranthaceae |
| Alfalfa | *Medicago sativa* | Fabaceae |
| Dried fat | *Elaeis guineensis/Brassica sp.* | Arecaceae/ Brassicaceae |
| "Lignobond" | *Picea abies, Pinus sylvestris* | Pinaceae |
| Limestone flour | | |
| Magnesium oxide | | |
| Monocalcium phosphate | | |
| Salt | | |
| Micro-minerals | | |
| Cu (Copper(II) sulphate) | | |
| Se (Sodium selenite) | | |
| Zn (Zinc sulphate) | | |
| Mn (Manganese(II) sulphate) | | |
| I (Calcium iodate) | | |
| Co (Cobalt (II) sulphate) | | |
| Vitamins | | |
| Vitamin A | | |
| Vitamin D3 | | |
| Vitamin E | | |

**Table S2:** Quantity of RF-80 feed consumed every day by the three reindeer (9/10, 10/10, 12/10) during the feeding experiment. Fresh feed was provided to the reindeer everyday *ad libitum* (i.e., 2,000 g). During the last three days of the experiment, the quantity of feed leftovers was estimated only visually.

| Date | Pellets eaten (g) | | |
|---|---|---|---|
| | **9/10** | **10/10** | **12/10** |
| 19.01.18 | 1445 | 751 | 1081 |
| 20.01.18 | 1316 | 831 | 1264 |
| 21.01.18 | 1437 | 1128 | 1468 |
| 22.01.18 | 850 | 860 | 848 |
| 23.01.18 | 787 | 572 | 204 |
| 24.01.18 | 1158 | 980 | 1026 |
| 25.01.18 | 1427 | 1140 | 1217 |
| 26.01.18 | 1485 | 972 | 1144 |
| 27.01.18 | 1499 | 1213 | 1425 |
| 28.01.18 | 1500 | 1212 | 1478 |
| 29.01.18 | 1481 | 915 | 1236 |
| 30.01.18 | 1470 | 879 | 1144 |
| 31.01.18 | 1486 | 984 | 1217 |
| 01.02.18 | 1498 | 1308 | 1418 |
| 02.02.18 | 1498 | 1453 | 1446 |
| 03.02.18 | 1000 | 858 | 930 |
| 04.02.18 | 1000 | 770 | 808 |
| 05.02.18 | 1000 | 846 | 874 |
| 06.02.18 | 1500 | 979 | 1242 |
| 07.02.18 | 1500 | 1155 | 1282 |
| 08.02.18 | 1500 | 1225 | 1283 |
| 09.02.18 | 1500 | 1201 | 1083 |
| 10.02.18 | 1500 | 1490 | 1470 |
| 11.02.18 | 1500 | 869 | 1322 |
| 12.02.18 | 1500 | 1101 | 862 |
| 13.02.18 | 1700 | 1196 | 1638 |
| 14.02.18 | 2000 | 1558 | 1708 |
| 15.02.18 | 2000 | 2000 | 2000 |
| 16.02.18 | 2000 | 2000 | 2000 |
| 17.02.18 | 2000 | 2000 | 2000 |

**Table S3:** Number of reindeer faeces collected immediately before, during and after the feeding experiment.

| Date | 9/10 | 10/10 | 12/10 |
|---|---|---|---|
| 22.01.18 | 3 | 1 | 1 |
| 24.01.18 | 4 | 3 | 4 |
| 25.01.18 | 4 | 4 | 4 |
| 26.01.18 | 4 | 4 | 4 |
| 27.01.18 | 4 | 4 | 4 |
| 28.01.18 | 4 | 4 | 4 |
| 29.01.18 | 4 | 4 | 4 |
| 30.01.18 | 4 | 4 | 4 |
| 31.01.18 | 4 | 5 | 4 |
| 01.02.18 | 4 | 4 | 4 |
| 02.02.18 | 4 | 3 | 4 |
| 03.02.18 | 2 | 2 | 2 |
| 04.02.18 | 2 | 2 | 2 |
| 05.02.18 | 2 | 2 | 2 |
| 06.02.18 | 2 | 2 | 2 |
| 07.02.18 | 2 | 2 | 2 |
| 08.02.18 | 2 | 2 | 2 |
| 09.02.18 | 2 | 2 | 2 |
| 10.02.18 | 2 | 2 | 3 |
| 11.02.18 | 1 | 1 | 1 |
| 12.02.18 | 2 | 2 | 2 |
| 13.02.18 | 2 | 2 | 2 |
| 14.02.18 | 2 | 2 | 2 |
| 15.02.18 | 1 | 1 | 1 |
| 16.02.18 | 1 | 1 | 1 |
| 17.02.18 | 1 | 1 | 1 |
| 18.02.18 | 1 | 1 | 1 |
| 19.02.18 | 1 | 1 | 1 |
| 20.02.18 | 1 | 1 | 1 |
| 21.02.18 | 1 | 1 | 1 |
| 22.02.18 | 1 | 1 | 1 |
| 02.03.18 | 1 | 1 | 2 |
| 06.03.18 | 1 | - | - |
| 09.03.18 | 1 | 1 | 1 |
| 13.03.18 | - | - | 1 |
| 16.03.18 | 1 | - | - |

| | | | |
|---|---|---|---|
| 20.03.18 | 1 | 1 | 1 |
| 23.03.18 | 1 | - | 1 |

**Table S4:** Details about the 234 faecal time-points selected for DNA metabarcoding diet analysis. In bold are 54 samples for which we could not distinguish PCR replicates due to erroneous tagging. NA correspond to sampling times that did not coincide with any of the birch or lichen feeding.

| Animal | Date | Sampling time (CET) | Time since birch feeding (hours) | Fed lichen (g) |
|---|---|---|---|---|
| 9/10 | 22.01.18 | 11:26 | -37 | NA |
| 9/10 | 24.01.18 | 06:10 | 6 | 20 |
| 9/10 | 24.01.18 | 13:30 | 13 | 20 |
| 9/10 | 24.01.18 | 22:48 | 23 | 20 |
| 9/10 | 25.01.18 | 03:19 | 27 | 20 |
| 9/10 | 25.01.18 | 08:10 | 32 | 20 |
| 9/10 | 25.01.18 | 13:01 | 37 | 20 |
| **9/10** | **25.01.18** | **21:21** | **45** | **20** |
| 9/10 | 26.01.18 | 03:21 | 51 | 20 |
| **9/10** | **26.01.18** | **09:00** | **57** | **20** |
| 9/10 | 26.01.18 | 16:38 | 64 | 20 |
| **9/10** | **26.01.18** | **22:00** | **70** | **20** |
| 9/10 | 27.01.18 | 06:00 | 78 | 20 |
| 9/10 | 27.01.18 | 08:50 | 81 | 20 |
| **9/10** | **27.01.18** | **14:35** | **86** | **20** |
| **9/10** | **27.01.18** | **22:00** | **94** | **20** |
| **9/10** | **28.01.18** | **06:00** | **102** | **20** |
| 9/10 | 28.01.18 | 11:54 | 108 | 20 |
| **9/10** | **28.01.18** | **16:43** | **113** | **20** |
| **9/10** | **28.01.18** | **22:00** | **118** | **20** |
| 9/10 | 29.01.18 | 06:00 | 126 | 500 |
| **9/10** | **29.01.18** | **09:13** | **129** | **500** |
| 9/10 | 29.01.18 | 14:14 | 134 | 500 |
| 9/10 | 29.01.18 | 23:59 | 144 | 500 |
| 9/10 | 30.01.18 | 03:22 | 147 | 500 |
| 9/10 | 30.01.18 | 07:44 | 152 | 500 |
| **9/10** | **30.01.18** | **14:30** | **158** | **500** |
| 9/10 | 30.01.18 | 23:55 | 168 | 500 |
| 9/10 | 31.01.18 | 02:45 | 171 | 500 |
| **9/10** | **31.01.18** | **11:58** | **180** | **500** |
| 9/10 | 31.01.18 | 14:31 | 182 | 500 |
| 9/10 | 31.01.18 | 16:12 | 184 | 500 |

| | | | | |
|---|---|---|---|---|
| 9/10 | 31.01.18 | 23:56 | 192 | 500 |
| 9/10 | 01.02.18 | 03:00 | 195 | 500 |
| 9/10 | 01.02.18 | 08:32 | 200 | 500 |
| **9/10** | **01.02.18** | **13:43** | **206** | **500** |
| 9/10 | 01.02.18 | 22:00 | 214 | 500 |
| **9/10** | **02.02.18** | **06:01** | **222** | **500** |
| 9/10 | 02.02.18 | 08:45 | 225 | 500 |
| 9/10 | 02.02.18 | 16:29 | 232 | 500 |
| 9/10 | 03.02.18 | 10:35 | 250 | 2 000 |
| 9/10 | 03.02.18 | 16:35 | 256 | 2 000 |
| 9/10 | 04.02.18 | 10:14 | 274 | 2 000 |
| 9/10 | 04.02.18 | 22:12 | 286 | 2 000 |
| 9/10 | 05.02.18 | 11:22 | 299 | 2 000 |
| 9/10 | 05.02.18 | 19:25 | 307 | 2 000 |
| 9/10 | 06.02.18 | 08:44 | 321 | 2 000 |
| 9/10 | 06.02.18 | 13:10 | 325 | 2 000 |
| 9/10 | 07.02.18 | 09:30 | 345 | 2 000 |
| 9/10 | 07.02.18 | 16:32 | 352 | 2 000 |
| 9/10 | 08.02.18 | 09:13 | 369 | NA |
| 9/10 | 08.02.18 | 19:19 | 379 | NA |
| 9/10 | 09.02.18 | 10:15 | 394 | NA |
| 9/10 | 09.02.18 | 19:26 | 403 | NA |
| **9/10** | **10.02.18** | **09:52** | **418** | **NA** |
| **9/10** | **10.02.18** | **19:15** | **427** | **NA** |
| 9/10 | 11.02.18 | 18:51 | 451 | NA |
| 9/10 | 12.02.18 | 07:59 | 464 | NA |
| 9/10 | 12.02.18 | 16:00 | 472 | NA |
| 9/10 | 13.02.18 | 09:39 | 489 | NA |
| **9/10** | **13.02.18** | **19:01** | **499** | **NA** |
| 9/10 | 14.02.18 | 08:01 | 512 | NA |
| **9/10** | **14.02.18** | **16:51** | **521** | **NA** |
| **9/10** | **15.02.18** | **08:57** | **537** | **NA** |
| **9/10** | **16.02.18** | **11:48** | **564** | **NA** |
| **9/10** | **17.02.18** | **10:48** | **587** | **NA** |
| **9/10** | **18.02.18** | **12:50** | **613** | **NA** |
| **9/10** | **19.02.18** | **12:52** | **637** | **NA** |
| **9/10** | **20.02.18** | **08:33** | **656** | **NA** |
| 9/10 | 21.02.18 | 12:59 | 685 | NA |
| 9/10 | 22.02.18 | 13:26 | 709 | NA |
| 9/10 | 02.03.18 | 15:25 | 903 | NA |
| **9/10** | **09.03.18** | **14:42** | **1071** | **NA** |
| 9/10 | 20.03.18 | 14:45 | 1335 | NA |
| **10/10** | **24.01.18** | **03:12** | **0** | **20** |

| | | | | |
|---|---|---|---|---|
| 10/10 | 24.01.18 | 08:16 | 5 | 20 |
| 10/10 | 24.01.18 | 17:00 | 14 | 20 |
| 10/10 | 24.01.18 | 22:27 | 19 | 20 |
| **10/10** | **25.01.18** | **03:15** | **24** | **20** |
| **10/10** | **25.01.18** | **11:00** | **32** | **20** |
| **10/10** | **25.01.18** | **17:10** | **38** | **20** |
| 10/10 | 25.01.18 | 23:17 | 44 | 20 |
| 10/10 | 26.01.18 | 03:25 | 48 | 20 |
| 10/10 | 26.01.18 | 11:05 | 56 | 20 |
| 10/10 | 26.01.18 | 16:34 | 61 | 20 |
| 10/10 | 26.01.18 | 22:00 | 67 | 20 |
| 10/10 | 27.01.18 | 06:00 | 75 | 20 |
| 10/10 | 27.01.18 | 09:00 | 78 | 20 |
| 10/10 | 27.01.18 | 16:06 | 85 | 20 |
| 10/10 | 27.01.18 | 23:59 | 93 | 20 |
| 10/10 | 28.01.18 | 06:00 | 99 | 20 |
| 10/10 | 28.01.18 | 11:32 | 104 | 20 |
| 10/10 | 28.01.18 | 16:37 | 109 | 20 |
| 10/10 | 28.01.18 | 00:00 | 93 | 20 |
| 10/10 | 29.01.18 | 06:00 | 123 | 20 |
| 10/10 | 29.01.18 | 10:38 | 127 | 20 |
| 10/10 | 29.01.18 | 16:25 | 133 | 20 |
| 10/10 | 29.01.18 | 23:57 | 141 | 20 |
| 10/10 | 30.01.18 | 03:00 | 144 | 500 |
| 10/10 | 30.01.18 | 11:58 | 153 | 500 |
| 10/10 | 30.01.18 | 14:21 | 155 | 500 |
| 10/10 | 30.01.18 | 23:55 | 165 | 500 |
| 10/10 | 31.01.18 | 05:56 | 171 | 500 |
| 10/10 | 31.01.18 | 11:55 | 177 | 500 |
| 10/10 | 31.01.18 | 16:07 | 181 | 500 |
| **10/10** | **31.01.18** | **23:58** | **189** | **500** |
| 10/10 | 01.02.18 | 03:01 | 192 | 500 |
| 10/10 | 01.02.18 | 09:21 | 198 | 500 |
| 10/10 | 01.02.18 | 16:28 | 205 | 500 |
| 10/10 | 01.02.18 | 23:57 | 213 | 500 |
| 10/10 | 02.02.18 | 03:01 | 216 | 500 |
| 10/10 | 02.02.18 | 08:42 | 222 | 500 |
| 10/10 | 02.02.18 | 16:24 | 229 | 500 |
| **10/10** | **02.02.18** | **19:50** | **233** | **500** |
| 10/10 | 03.02.18 | 11:34 | 248 | 2 000 |
| 10/10 | 03.02.18 | 20:02 | 257 | 2 000 |
| **10/10** | **04.02.18** | **10:31** | **271** | **2 000** |
| 10/10 | 04.02.18 | 22:14 | 283 | 2 000 |

| | | | | |
|---|---|---|---|---|
| **10/10** | **05.02.18** | **11:16** | **296** | **2 000** |
| **10/10** | **05.02.18** | **19:23** | **304** | **2 000** |
| **10/10** | **06.02.18** | **08:42** | **318** | **2 000** |
| **10/10** | **06.02.18** | **19:28** | **328** | **2 000** |
| **10/10** | **07.02.18** | **09:28** | **342** | **2 000** |
| 10/10 | 07.02.18 | 19:25 | 352 | 2 000 |
| 10/10 | 08.02.18 | 09:30 | 366 | NA |
| **10/10** | **08.02.18** | **19:15** | **376** | **NA** |
| 10/10 | 09.02.18 | 11:10 | 392 | NA |
| 10/10 | 09.02.18 | 19:00 | 400 | NA |
| **10/10** | **10.02.18** | **10:13** | **415** | **NA** |
| 10/10 | 10.02.18 | 19:06 | 424 | NA |
| 10/10 | 11.02.18 | 18:58 | 448 | NA |
| **10/10** | **12.02.18** | **09:17** | **462** | **NA** |
| 10/10 | 12.02.18 | 18:58 | 472 | NA |
| 10/10 | 13.02.18 | 11:55 | 489 | NA |
| 10/10 | 13.02.18 | 19:05 | 496 | NA |
| **10/10** | **14.02.18** | **09:24** | **510** | **NA** |
| 10/10 | 14.02.18 | 12:50 | 514 | NA |
| 10/10 | 15.02.18 | 10:23 | 535 | NA |
| 10/10 | 16.02.18 | 09:51 | 559 | NA |
| 10/10 | 17.02.18 | 09:53 | 583 | NA |
| **10/10** | **18.02.18** | **09:56** | **607** | **NA** |
| 10/10 | 19.02.18 | 12:51 | 634 | NA |
| **10/10** | **20.02.18** | **08:16** | **653** | **NA** |
| 10/10 | 21.02.18 | 13:22 | 682 | NA |
| **10/10** | **22.02.18** | **12:33** | **705** | **NA** |
| **10/10** | **22.01.18** | **14:15** | **-37** | **NA** |
| **10/10** | **22.01.18** | **16:00** | **-35** | **NA** |
| 10/10 | 22.01.18 | 19:33 | -31 | NA |
| 10/10 | 02.03.18 | 14:55 | 900 | NA |
| 10/10 | 06.03.18 | 15:20 | 996 | NA |
| 10/10 | 09.03.18 | 15:08 | 1068 | NA |
| 10/10 | 16.03.18 | 14:50 | 1236 | NA |
| 10/10 | 20.03.18 | 14:38 | 1331 | NA |
| 10/10 | 23.03.18 | 14:22 | 1403 | NA |
| 12/10 | 22.01.18 | 14:00 | -35 | NA |
| **12/10** | **24.01.18** | **01:17** | **0** | **20** |
| 12/10 | 24.01.18 | 11:10 | 10 | 20 |
| 12/10 | 24.01.18 | 13:30 | 12 | 20 |
| 12/10 | 24.01.18 | 22:28 | 21 | 20 |
| 12/10 | 25.01.18 | 03:08 | 26 | 20 |
| 12/10 | 25.01.18 | 11:00 | 34 | 20 |

| | | | | |
|---|---|---|---|---|
| 12/10 | 25.01.18 | 15:50 | 39 | 20 |
| **12/10** | **25.01.18** | **21:13** | **44** | **20** |
| 12/10 | 25.01.18 | 23:13 | 46 | 20 |
| 12/10 | 26.01.18 | 05:59 | 53 | 20 |
| 12/10 | 26.01.18 | 11:07 | 58 | 20 |
| 12/10 | 26.01.18 | 13:24 | 60 | 20 |
| 12/10 | 26.01.18 | 22:00 | 69 | 20 |
| 12/10 | 27.01.18 | 06:00 | 77 | 20 |
| 12/10 | 27.01.18 | 08:50 | 80 | 20 |
| 12/10 | 27.01.18 | 14:30 | 85 | 20 |
| 12/10 | 27.01.18 | 23:59 | 95 | 20 |
| 12/10 | 28.01.18 | 06:00 | 101 | 20 |
| 12/10 | 28.01.18 | 11:55 | 107 | 20 |
| 12/10 | 28.01.18 | 16:33 | 111 | 20 |
| 12/10 | 28.01.18 | 23:59 | 119 | 20 |
| 12/10 | 29.01.18 | 06:00 | 125 | 500 |
| 12/10 | 29.01.18 | 10:42 | 129 | 500 |
| 12/10 | 29.01.18 | 14:07 | 133 | 500 |
| 12/10 | 29.01.18 | 23:50 | 143 | 500 |
| **12/10** | **30.01.18** | **03:00** | **146** | **500** |
| 12/10 | 30.01.18 | 11:41 | 154 | 500 |
| 12/10 | 30.01.18 | 14:11 | 157 | 500 |
| **12/10** | **30.01.18** | **23:55** | **167** | **500** |
| 12/10 | 31.01.18 | 05:50 | 173 | 500 |
| 12/10 | 31.01.18 | 11:07 | 178 | 500 |
| 12/10 | 31.01.18 | 14:23 | 181 | 500 |
| 12/10 | 31.01.18 | 21:57 | 189 | 500 |
| 12/10 | 01.02.18 | 02:54 | 194 | 500 |
| 12/10 | 01.02.18 | 09:24 | 200 | 500 |
| 12/10 | 01.02.18 | 16:24 | 207 | 500 |
| 12/10 | 01.02.18 | 22:21 | 213 | 500 |
| 12/10 | 02.02.18 | 02:56 | 218 | 500 |
| 12/10 | 02.02.18 | 10:10 | 225 | 500 |
| 12/10 | 02.02.18 | 16:20 | 231 | 500 |
| **12/10** | **02.02.18** | **19:46** | **234** | **500** |
| 12/10 | 03.02.18 | 10:24 | 249 | 2 000 |
| 12/10 | 03.02.18 | 17:40 | 256 | 2 000 |
| 12/10 | 04.02.18 | 10:14 | 273 | 2 000 |
| 12/10 | 04.02.18 | 22:09 | 285 | 2 000 |
| 12/10 | 05.02.18 | 11:14 | 298 | 2 000 |
| 12/10 | 05.02.18 | 16:02 | 303 | 2 000 |
| 12/10 | 06.02.18 | 10:40 | 321 | 2 000 |
| 12/10 | 06.02.18 | 19:26 | 330 | 2 000 |

| | | | | |
|---|---|---|---|---|
| **12/10** | **07.02.18** | **09:43** | **344** | **2 000** |
| 12/10 | 07.02.18 | 16:19 | 351 | 2 000 |
| 12/10 | 08.02.18 | 08:23 | 367 | NA |
| 12/10 | 08.02.18 | 19:10 | 378 | NA |
| 12/10 | 09.02.18 | 10:30 | 393 | NA |
| 12/10 | 09.02.18 | 19:28 | 402 | NA |
| 12/10 | 10.02.18 | 09:11 | 416 | NA |
| 12/10 | 10.02.18 | 10:17 | 417 | NA |
| **12/10** | **10.02.18** | **19:06** | **426** | **NA** |
| **12/10** | **11.02.18** | **18:51** | **450** | **NA** |
| 12/10 | 12.02.18 | 09:17 | 464 | NA |
| 12/10 | 12.02.18 | 16:00 | 471 | NA |
| 12/10 | 13.02.18 | 11:55 | 491 | NA |
| 12/10 | 13.02.18 | 19:07 | 498 | NA |
| 12/10 | 14.02.18 | 08:41 | 511 | NA |
| 12/10 | 14.02.18 | 16:38 | 519 | NA |
| 12/10 | 15.02.18 | 09:19 | 536 | NA |
| 12/10 | 16.02.18 | 11:21 | 562 | NA |
| 12/10 | 17.02.18 | 10:28 | 585 | NA |
| 12/10 | 18.02.18 | 09:56 | 609 | NA |
| 12/10 | 19.02.18 | 12:52 | 636 | NA |
| 12/10 | 20.02.18 | 09:14 | 656 | NA |
| 12/10 | 21.02.18 | 12:29 | 683 | NA |
| 12/10 | 22.02.18 | 13:03 | 708 | NA |
| 12/10 | 02.03.18 | 14:50 | 902 | NA |
| 12/10 | 02.03.18 | 14:57 | 902 | NA |
| 12/10 | 09.03.18 | 14:47 | 1070 | NA |
| 12/10 | 13.03.18 | 15:50 | 1167 | NA |
| 12/10 | 20.03.18 | 14:41 | 1333 | NA |
| 12/10 | 23.03.18 | 14:10 | 1405 | NA |

**Table S5:** Details on the two DNA metabarcoding primer sets used for the analysis of reindeer diets. Abbreviations: bp = base pair.

| Taxonomic group | Spermatophyta (seed plants) | Eukaryota |
|---|---|---|
| Primer name | *Sper01* | *Euka02* |
| Target gene | trn*L* (UAA) | 18S V7 region |
| Sequence (5'-3') | GGGCAATCCTGAGCCAA<br>CCATTGAGTCTCTGCACCTATC | TTTGTCTGSTTAATTSCG<br>CACAGACCTGTTATTGC |
| Size range (bp) | 10 – 220 (average: 48) | 36 – 892 (average: 123) |
| Annealing temperature | 52°C | 45°C |
| Reference | Taberlet et al. 2007[1] | Guardiola et al. 2015[2] |

[1]Taberlet P, Coissac E, Pompanon F, Gielly L, Miquel C, Valentini A, Vermat T, Corthier G, Brochmann C, Willerslev E (2007) Power and limitations of the chloroplast trnL (UAA) intron for plant DNA barcoding. Nucleic Acids Research 35(3): e14. doi: 10.1093/nar/gkl938; [2]Guardiola M, Uriz MJ, Taberlet P, Coissac E, Wangensteen OS, Turon X (2015) Deep-Sea, Deep-Sequencing: Metabarcoding Extracellular DNA from Sediments of Marine Canyons. PLoS ONE 10(10): e0139633. https://doi.org/10.1371/journal.pone.0139633

**Table S6:** Summary of the six synthetic DNA standards used as a mock community PCR positive control with the Sper01 plant primers.

| Synthetic standard | Dilution | Reference sequence |
|---|---|---|
| SS_01_Sper01 | 1 | taagtctcgcactagttgtgacctaacgaatagagaattctataagacgtgttgtcccat |
| SS_02_ Sper01 | 1/2 | gtgtatggtatatttgaataatattaaatagaatttaatcaatctttacatcgcttaata |
| SS_03_ Sper01 | 1/4 | cacaatgctcggtaactagaagcatttgta |
| SS_04_ Sper01 | 1/8 | attgaatgaaaagattattcgatatagaat |
| SS_05_ Sper01 | 1/16 | agaacgctagaatctaagatggggggggggatgagtaagatatttatcagtaacatatga |
| SS_06_ Sper01 | 1/32 | atttttgtaactcattaacaattttttttttgatgtatcataagtactaaactagttact |
| SS_01_Euka02 | 1 | taagtctcgcactagttgtgacctaacgaatagagaattctataagacgtgttgtcccat |

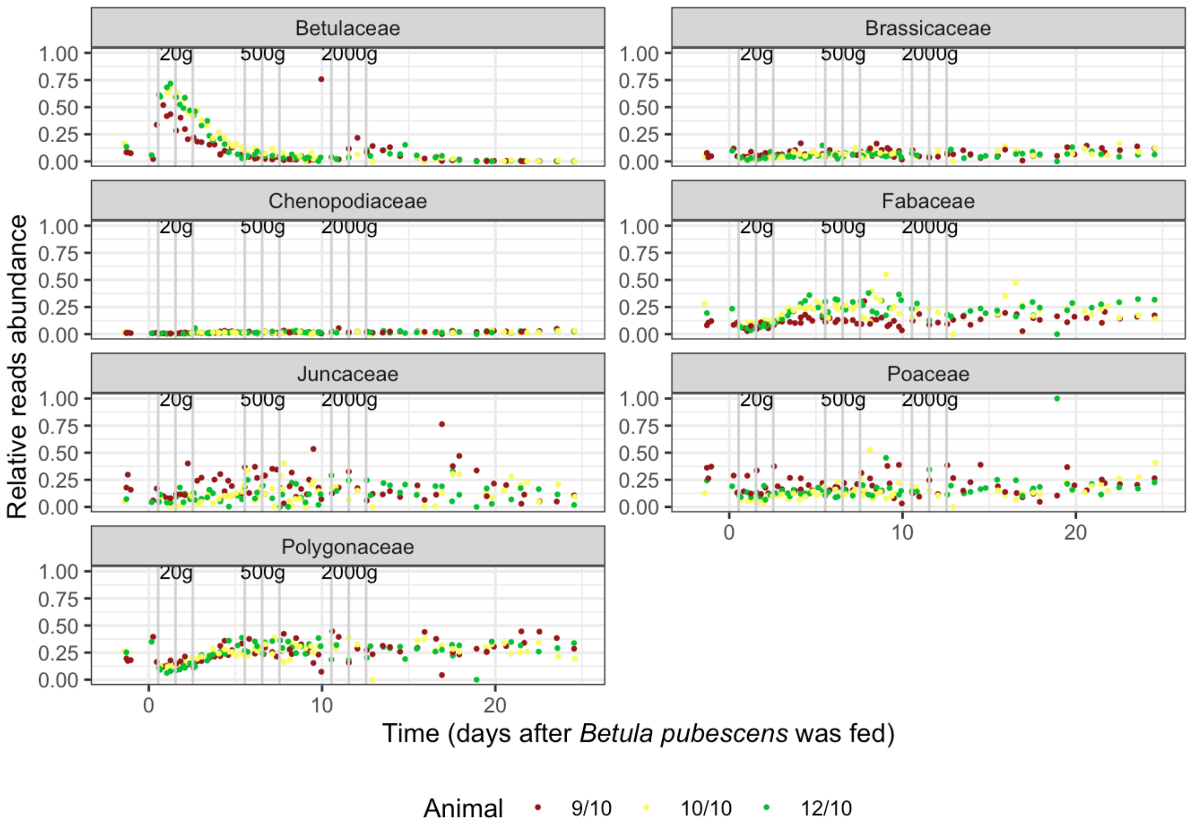

**Figure S1:** Overview of the seed plant families (and the change in their relative proportions over time), as detected from the reindeer faeces with the *Sper01* primer set. The three sets of vertical lines correspond to the lichen feeding events (20, 500 and 2,000 g of lichen biomass, respectively). The dotted vertical line marks the end of the experiment.

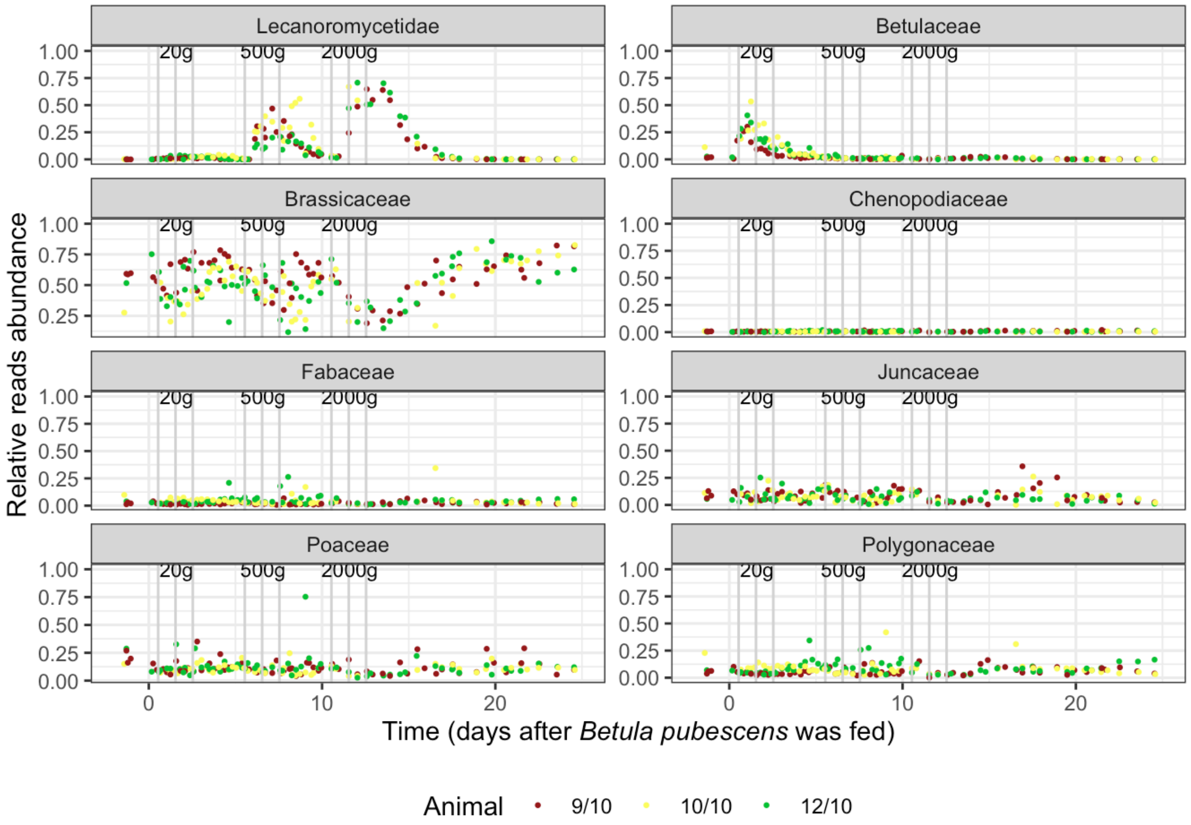

**Figure S2:** Overview of the seed plant families and lichen (subclass Lecanoromycetidae) (and the change in their relative proportions over time), as detected from the reindeer faeces with the *Euka02* primer set. The three sets of vertical lines correspond to the lichen feeding events (20, 500 and 2,000 g of lichen biomass, respectively). The dotted vertical line marks the end of the experiment.

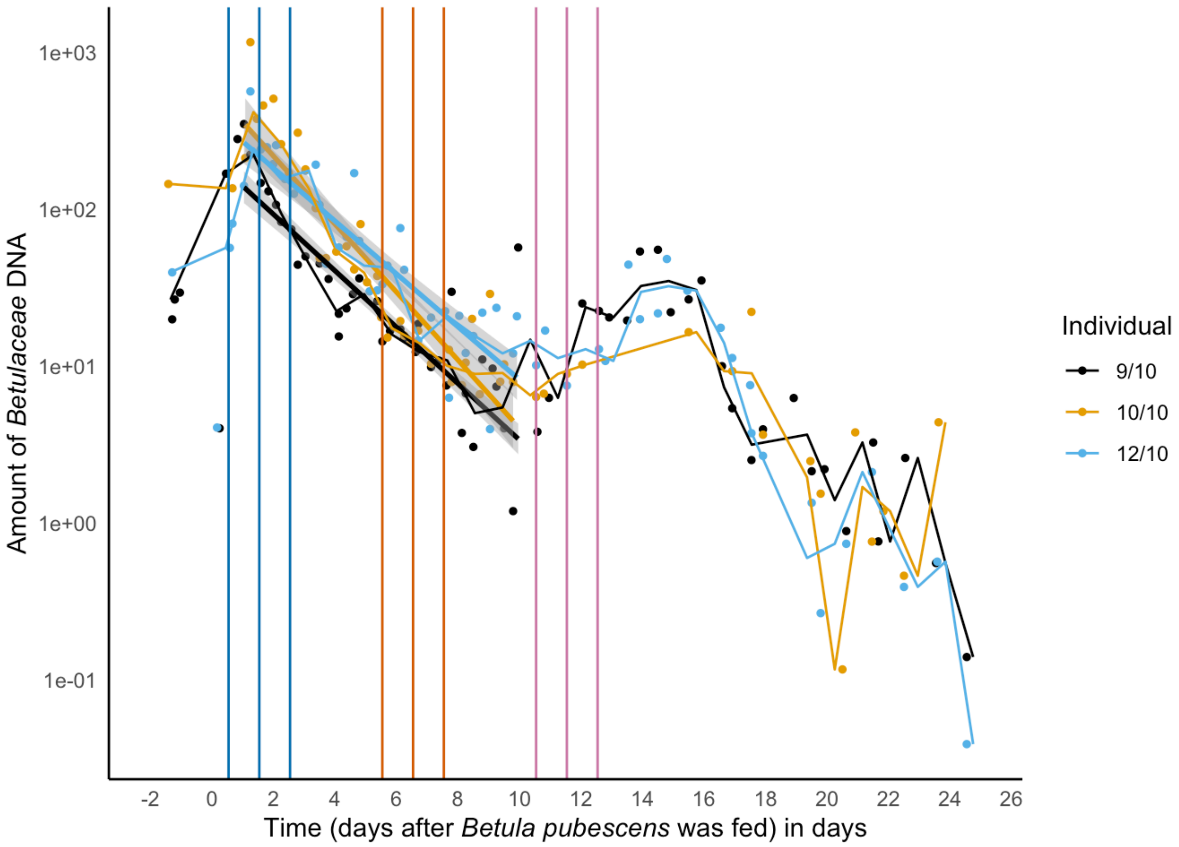

**Figure S3.** Proportion of birch DNA (*Betula pubescence*) in reindeer faecal samples, as detected with the plant Sper01 primer set. Solid lines represent the linear regression whose slope was used to estimate DNA half-life detectability for each individual reindeer. Time 0 corresponds to the time birch was introduced to the reindeer (samples collected prior to time 0 are the reference faecal samples before the experiment). Colored vertical lines correspond to the recurrent lichen feeding events.
